# Myeloid Dnmt3a loss disrupts macrophage polarization and leads to an age-dependent impairment of hematopoietic homeostasis

**DOI:** 10.64898/2026.11.17.766737

**Authors:** Dora Bordoni, Celia Escudero-Hernández, Franziska Kimmig, Neha Mishra, Antonella Fazio, Florian Tran, Simon Imm, Finn Hinrichsen, Saskia Weber-Stiehl, Stephanie Stengel, Felix Sommer, Maren Falk-Paulsen, Philip Rosenstiel

## Abstract

Genetic variants in the de novo DNA methyltransferase DNMT3A are associated with inflammatory bowel disease (IBD), but its contribution to intestinal immune homeostasis remains poorly understood. Here we show that myeloid Dnmt3a deletion disrupts macrophage polarization and promotes inflammatory and phagocytic responses. Using mice with LysM-driven Dnmt3a deletion, we combine methylation and transcriptomic profiling with functional characterization of bone marrow-derived macrophages and experimental colitis. Dnmt3a expression increases during M2 polarization and decreases following inflammatory stimulation. Despite IL-4/IL-13 stimulation, Dnmt3a-deficient macrophages exhibit reduced M2 marker expression and increased expression of inflammatory genes, including Irg1 and Tnf. These cells acquire M1-like morphological and functional features and impair regeneration of wounded intestinal epithelial monolayers in co-culture. The phenotype is accentuated in macrophages from mice older than 34 weeks, with enhanced phagocytosis, reduced arginase activity and increased inducible nitric oxide synthase activity. In vivo, young mutant mice show increased splenic T-cell numbers, fewer macrophages but only subtle changes in susceptibility to dextran sodium sulfate-induced experimental colitis. Older mice develop altered bone marrow architecture, enhanced myeloproliferation and spontaneous hepatosplenomegaly. These findings establish DNMT3A as a regulator of macrophage polarization and identify a myeloid mechanism through which its loss may contribute to IBD pathogenesis.

**SYNOPSIS:** The *de novo* DNA methyltransferase *DNMT3A* is required for physiological macrophage polarization as its depletion leads to a pro-inflammatory skewing, especially of M2 bone-marrow derived macrophage (BMDM) polarization with IL-4 and IL-13. Myeloid lack of *Dnmt3a* promotes hepatosplenomegaly by HSC hyperproliferation.

**KEY SUMMARY:**

1. Summary of the established knowledge on this subject:
  - Genetic variants in the *de novo* DNA methyltransferase *DNMT3A* associate with increased risk for inflammatory bowel disease (IBD).
  - Macrophages are key gatekeepers for intestinal immune homeostasis, and molecular pathways governing monocyte-to-macrophage differentiation during inflammation remain poorly understood.
2. What are the significant and/or new findings of this study?
  - Normal *DNMT3A* function is required for physiological macrophage polarization.
  - Bone-marrow derived macrophages (BMDM) lacking *Dnmt3a* expression lead to a pro-inflammatory skewing, particularly after polarization with M2-inducing cytokines IL-4 and IL-13.
  - While mice lacking *Dnmt3a* myeloid expression display only subtle changes in experimental colitis compared to littermate controls, they develop hepatosplenomegaly with age.
  - Understanding perturbed epigenetic imprints on macrophage polarization in IBD could provide novel disease-progression biomarkers and highlight new druggable targets.

## INTRODUCTION

Intestinal macrophages are key, versatile antigen presenting cells that contribute to immune homeostasis in the gut. These originate from migrated circulating inflammatory monocytes that were attracted by low-grade interferon (IFN)-driven inflammation induced by the microbiota which, in turn, is controlled by an IL-10 anti-inflammatory response^1–3^. Macrophages are naturally programmed to rapidly respond to challenges while avoiding sustained hyperactivation of proinflammatory features, e.g. cytokines or metalloproteinases (MMP) that could damage the tissue^4^. Depending on the microenvironment, macrophages are thought to polarize into two main functional phenotypes comprising activated M1 catabolic pro-glycolytic, proinflammatory macrophages involved in antimicrobial host defense (after priming with IFN-γ and lipopolysaccharide, LPS) and alternatively activated M2 anabolic, anti-inflammatory macrophages that counteract inflammation and stimulate tissue repair. M2 polarization can typically be induced by stimulation with type II interleukins IL-4 and IL-13)^5,6^. The simplistic concept has been significantly challenged in the past years by scRNAseq-based studies showing heterogeneity of cell populations beyond this simple dichotomy^7–9^. Despite this complexity, the importance of sustaining a well-balanced ratio of macrophage phenotypes as a determinant of immune homeostasis is overt and can be considered an important target for therapeutic interference with the immune system. As such, a perturbed equilibrium of macrophages has been shown to be part of sustained low-grade inflammation (e.g. during ageing, so-called “inflammaging”) as well as chronic inflammatory processes such as IBD^10,11^. However, the exact mechanisms of how macrophages are re-programmed to fuel chronic inflammation are still unclear. Epigenetic modifications have been implicated in macrophage cellular plasticity^4^. In the context of intestinal inflammation, histone deacetylase HDAC3 inhibition via butyrate has been suggested to modulate selective macrophage functions toward antimicrobial host defense, while avoiding tissue-destructive inflammation (i.e. downregulating IL-10 and IL-12 expression)^10,12,13^. In IBD, polymorphisms in the gene coding for the de novo methyltransferase *DNMT3A* have been associated with increased risk for the disease^14^. In addition to a SNP-dependent eQTL effect of the IBD associated variants, which leads to downregulation of DNMT3A transcript levels, inflammatory activity in IBD is characterized by lower levels of *DNMT3A* in intestinal epithelial cells^15,16^. Our previous study demonstrated barrier defects when *DNMT3A* was ablated in intestinal epithelial cells in mice, which was linked to increased susceptibility to dextran sodium sulfate (DSS)-induced colitis^15^. However, an overall hypomethylation, especially on immune-related genes, has been demonstrated in the intestinal mucosa of IBD patients^17– 19^, suggesting deregulation of DNA methylation also in other cell types.

Here, we investigated the effects of myeloid Dnmt3a deletion in mice by integrating methylation and transcriptomic profiling with functional analyses on bone-marrow derived macrophages and *in vivo* phenotyping. We examined macrophage responses to defined polarizing stimuli, their effects on intestinal epithelial gap closure and barrier formation, and outcomes in experimental colitis and older mice.**RESULTS**

### *Dnmt3a* levels are upregulated during M2 macrophage polarization and ablation leads to skewed levels of M1/M2-associated features

To elucidate the role of *Dnmt3a* for macrophage differentiation, we generated C57/BL6 mice lacking *Dnmt3a* in the myelomonocytic lineage (Dnmt3a^ΔLysM^) using a conditional Dnmt3a allele with the exons 13–17 flanked by loxP sites^15^ crossed with LysM mice expressing Cre recombinase under the control of the Lyz2 promoter (Suppl. fig. 1). While both intestinal lamina propria (LP) CD11b+ macrophages and bone-marrow derived macrophages (BMDM) lacked *Dnmt3a* expression in Dnmt3a^ΔLysM^ mice (Suppl. fig. 1A-G), *in vitro* proportions of CD11b^+^ cells differentiated from the bone marrow (BMDM) remained constant (Suppl. fig. 1H-I).

Next, we examined the impact of *in vitro* polarization of BMDMs on the expression of *Dnmt3a*. To induce M1 and M2 phenotypes, cells were stimulated for 96 hours with LPS and IFN-γ or with IL-4 and IL-13, respectively, following an established protocol. In BMDMs from WT mice, Dnmt3a protein levels increased significantly in M2-polarized macrophages (days 6-9) while they remained almost undetectable during M1 polarization and in resting cells (Figure 1A-B), which mirrors earlier findings in Th1/2 differentiation^20^.

**Figure 1:**
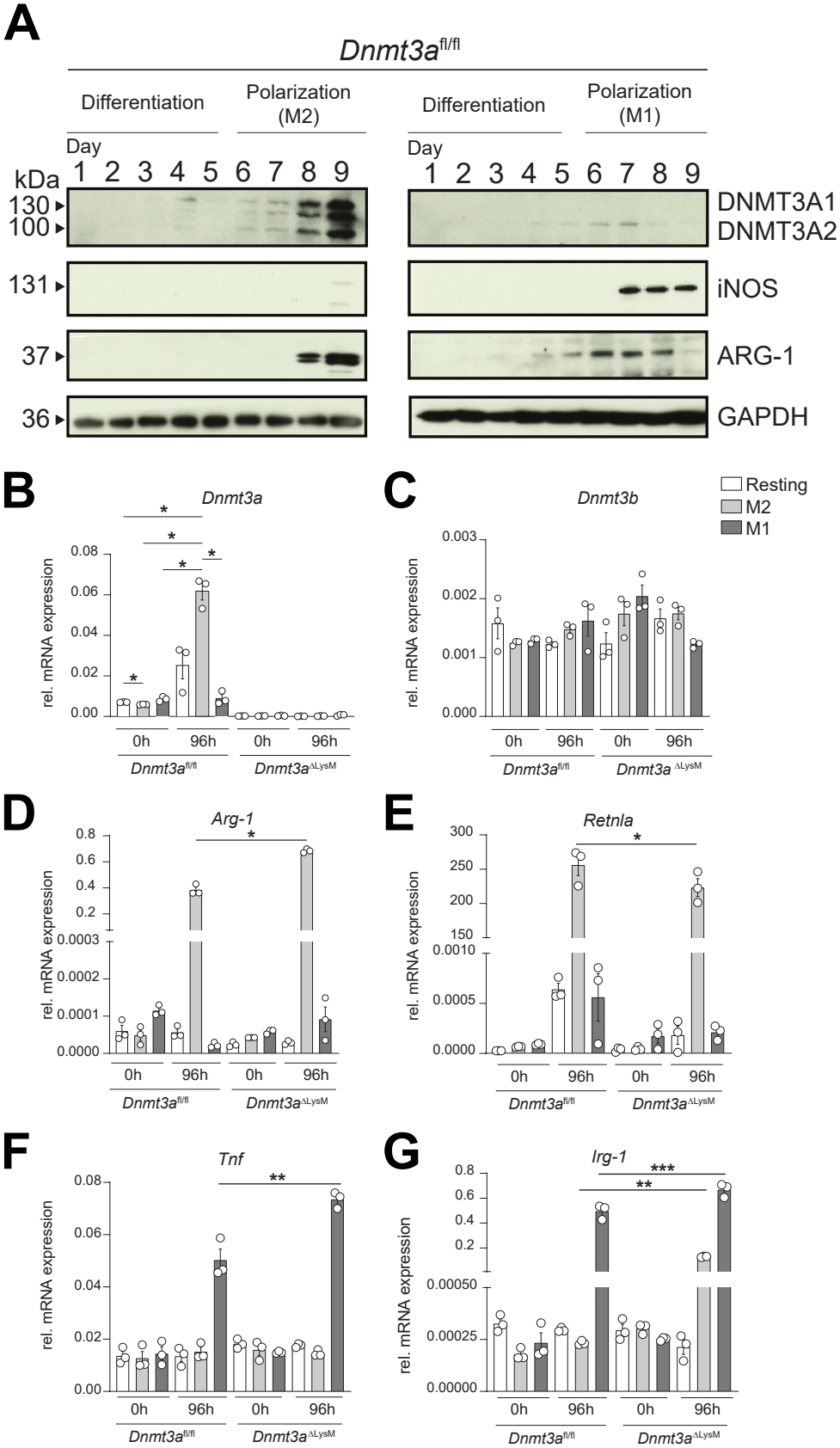
Lack of *Dnmt3a* leads to impaired macrophage polarization. (A) Dnmt3a, iNOS and Arg-1 progressive protein detection during *in vitro* BMDM 5-day differentiation with 20 ng/mL m-CSF and subsequent 4-day polarization into M1/proinflammatory iNOS^+^ and M2/anti-inflammatory Arg-1^+^ phenotypes from Dnmt3a^fl/fl^ mice. Gapdh was used as a reference. (B-G) Relative gene expression of *Dnmt3a* (B), *Dnmt3b* (C), *Arg-1* (D), *Retnla* (E), *Tnf* (F) and *Irg-1* (G) transcripts on *in vitro* polarized BMDMs from Dnmt3a^fl/fl^ and Dnmt3a^ΔLysM^ mice. In all cases, BMDM were polarized using 20 ng/mL m-CSF for the resting program, 100 ng/mL LPS and 50 ng/mL IFN-g for the M1 program, and 50 ng/mL IL-4 and 50 ng/mL IL-13 for the M2 program. Arg-1, Arginase 1; Irg-1, immune-responsive gene 1; m-CSF, macrophage colony stimulating factor; Tnf, tumor necrosis factor; Retnla, resistin like alpha. All results are based on n = 3-6 biological replicates per group. Resting BMDM samples are shown in white, M2 BMDM samples are shown in grey, and M1 BMDM samples in dark grey. Data are represented as mean ± SEM. Statistically significant differences are shown as *p<0,05, **p<0,001, ***p<0,0002 ****, p<0,0001.

To investigate the effect of Dnmt3a ablation on *in vitro* polarization, we tested selected mRNA markers indicative of M1 or M2 differentiation comparing WT to Dnmt3a^ΔLysM^-derived cells. M1 markers *Tnf* and *Irg1* mRNA expression were increased in M1-polarized macrophages from Dnmt3a^ΔLysM^ mice compared to their Dnmt3a^fl/fl^ counterparts (Figure 1B-G). M2 polarized Dnmt3a^ΔLysM^ cells not only failed to achieve a full M2 polarization according to decreased resistin-like molecule alpha ( *Retnla*) mRNA expression, but also showed a tendency towards M1 polarization according to increased *Tnf* and *Irg1* mRNA expression (Figure 1B-G) The latter gene encodes the metabolic enzyme aconitate decarboxylase (ACOD1), known to suppress M2 polarization by elevating itaconate^21,22^.

Together, these findings suggest that ablation of Dnmt3a leads to a complex perturbation of *in vitro* macrophage polarization with a potential increase of pro-inflammatory features in Dnmt3a^ΔLysM^ derived M1 and M2 cells.

### Dnmt3a deletion induces hypomethylation and transcriptional dysregulation in macrophages that could lead to morphological alterations, proinflammatory responses, and malignant transformation

To gain a systematic insight into skewed polarization patterns of Dnmt3a^ΔLysM^ macrophages, we explored methylation and transcriptional differences between resting, M1 and M2 polarized BMDMs using epigenome-wide methylation array and RNAseq analyses. Since Dnmt3a loss-of-function mutations increase hematopoietic stem cell (HSC) proliferation and are associated with age-dependent clonal hematopoiesis^23–25^, we utilized BMDMs from young (8-10 weeks old) and old (>34 weeks old) mice (Figures 2-3, Suppl. fig. 2-3).

**Figure 2.**
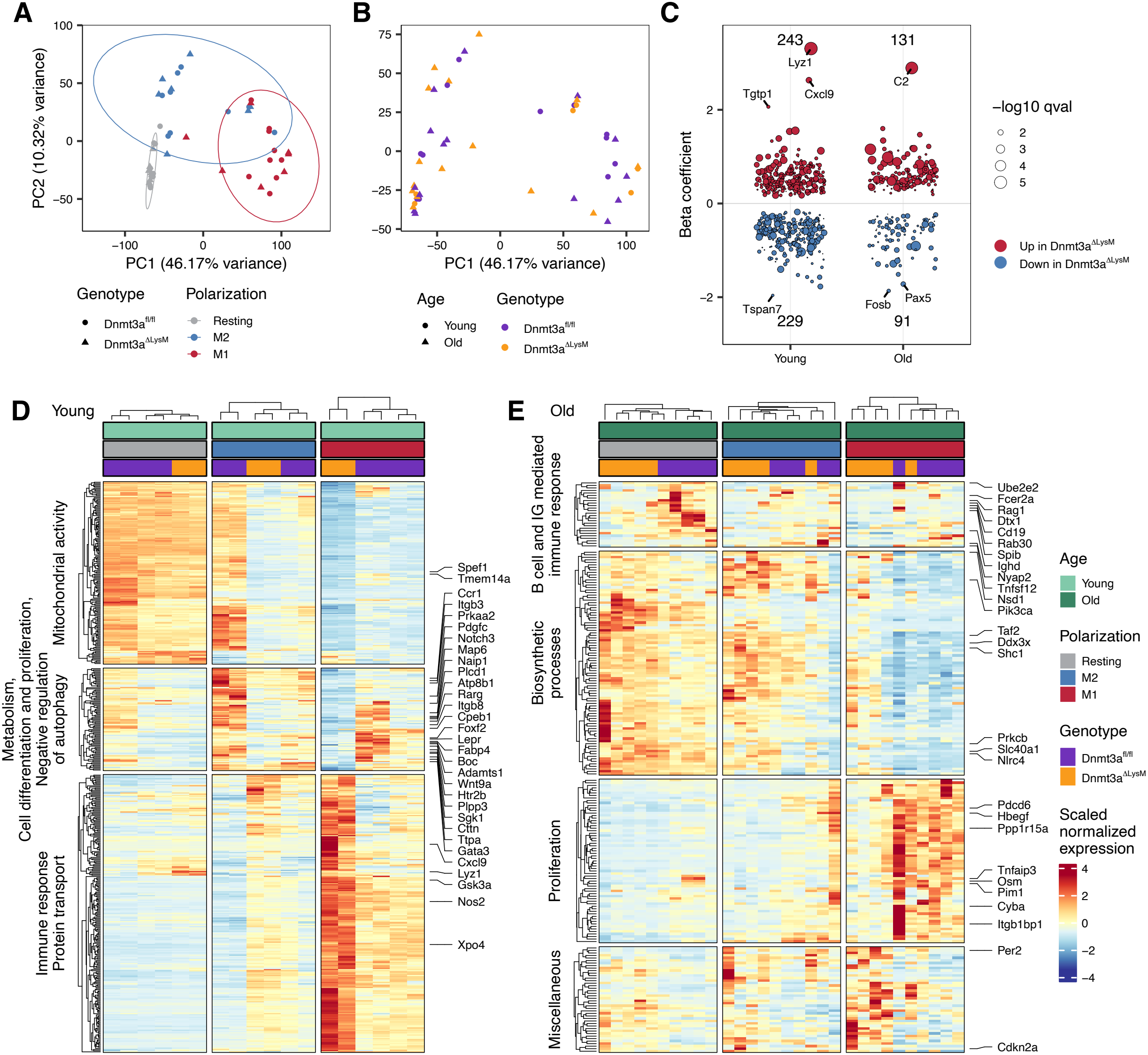
Dnmt3a depletion induces transcriptional cytoskeletal and proinflammatory changes in M2-polarized BMDMs from Dnmt3a^ΔLysM^mice. (A-B) Principal component analysis (PCA) of BMDM transcriptome from young (8-10 weeks old) and aged (>34 weeks-old) Dnmt3a^fl/fl^ and Dnmt3a^ΔLysM^ mice. (C) Mixed linear model beta coefficient (accounting for polarization and genotype) across different age groups displaying upregulated differentially expressed genes (DEGs) in red and downregulated DEGs in blue. Dot size corresponds to the FDR adjusted p-value. (D-E) Heatmap of DEGs showing the scaled normalized expression in resting, M1 and M2 polarized BMDMs from young (D) and aged (E) Dnmt3a^fl/fl^ and Dnmt3a^ΔLysM^ mice. DEG clusters are manually annotated and selected genes labelled. Analyses shown in A-E were conducted on n=4 young Dnmt3a^fl/fl^, n=2 young Dnmt3a^ΔLysM^, n=5 aged Dnmt3a^fl/fl^, and n=5 aged Dnmt3a^ΔLysM^ mice, with one BMDM sample per polarization per mouse; total n=48.

**Figure 3.**
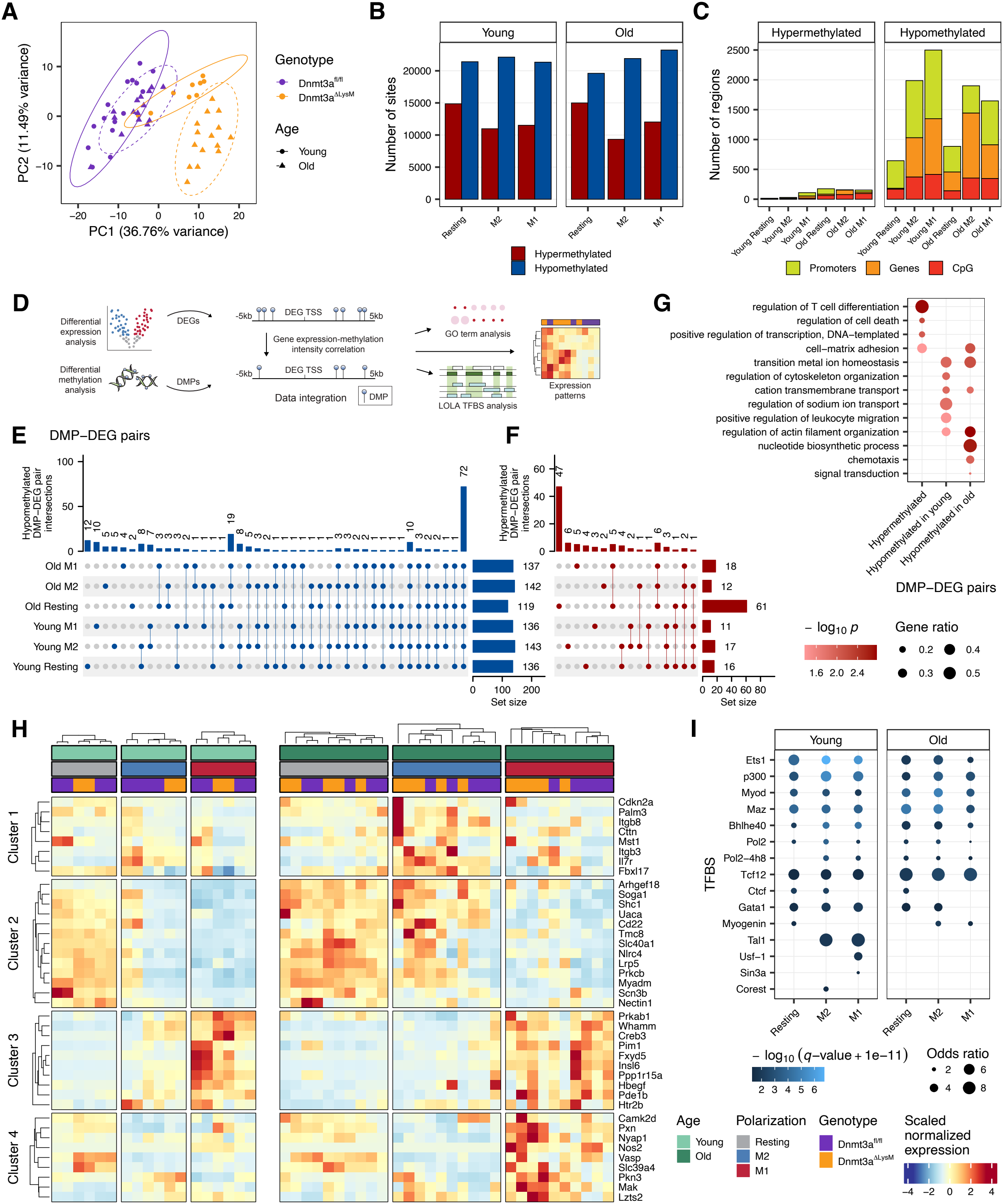
Dnmt3a depletion induces transcriptional cytoskeletal, proinflammatory and malignant changes in M2-polarized BMDMs from Dnmt3a^ΔLysM^mice. (A) Principal component analysis (PCA) of the DNA methylation array data for BMDMs from young (8-10 weeks old) and aged (>34 weeks-old) Dnmt3a^fl/fl^ and Dnmt3a^ΔLysM^ mice according to polarization and genotype. (B) Number of hypermethylated (red) or hypomethylated (blue) sites per BMDM group. (C) Number of promoter (yellow), genes (orange) or CpG island (red) regions that were found hypo- or hypermethylated. (D) Workflow for the integration of DNA methylation and transcriptomic analyses. (E-F) Intersection plots of differentially methylated positions (DMP) correlated with differentially expressed genes (DEG) in BMDM samples from young and aged Dnmt3a^fl/fl^ and Dnmt3a^DLysM^ mice according to hypomethylation (E) or hypermethylation (F). (G) Gene ontology biological processes enriched in DMP-DEG pairs in polarized BMDMs from young and aged Dnmt3a^DLysM^ mice compared to Dnmt3a^fl/fl^ mice. Dot sizes indicate gene ratio as the ratio of significant DEGs to annotated genes in each gene set term and colours corresponds to the p-value of enrichment. (H) Heatmaps showing scaled normalized expression of the leading DMP-DEG pairs between Dnmt3a^fl/fl^ and Dnmt3a^DLysM^ samples from young and aged mice (n=4 young Dnmt3a^fl/fl^, n=2 young Dnmt3a^DLysM^, n=5 aged Dnmt3a^fl/fl^, and n=5 aged Dnmt3a^DLysM^ mice, with one BMDM sample per polarization per mouse; total n=48) enriched in various biological processes. (I) Enrichment of transcription factor binding sites (TFBS) in DMP-DEG pairs that display hypomethylation or hypermethylation in polarized BMDMs from young and aged Dnmt3a^DLysM^ mice. Analyses shown in A-C were conducted on n=5 young Dnmt3a^fl/fl^, n=3 young Dnmt3a^DLysM^, n=5 aged Dnmt3a^fl/fl^, and n=5 aged Dnmt3a^DLysM^ mice, with one BMDM sample per polarization per mouse; total n=54. Samples from young (8-10 weeks-old) or old (>34 weeks-old) mice are shown in light or dark green, respectively; while resting, M1 and M2 polarized BMDMs are shown in grey, red or blue, respectively; and wild-type (Dnmt3a^fl/fl^) or knockout (Dnmt3a^DLysM^) are shown in purple or orange. Heatmap columns are split according to hierarchical clustering and named according to key terms.

PCA from RNA-seq data separated samples according to BMDM polarization status; however, genotype or age failed to separate the different groups (Figure 2A-B). Using polarization and genotype as fixed effects, we identified 472 differentially expressed genes (DEGs) in BMDMs from young mice. Among these, genes associated with mitochondrial function, metabolism, cell differentiation, proliferation, and morphogenesis were downregulated in both M1 and M2 Dnmt3a^ΔLysM^ BMDMs compared to their WT counterparts, while DEGs related to immune pathways and protein transport were increased (Figure 2C-D, Suppl. Fig. 2A-B). On the other hand, the 222 DEGs that differed in BMDMs from aged mice pointed towards an increased biosynthesis and decreased Ig-mediated immune responses and proliferation in M2 BMDMs from Dnmt3a^ΔLysM^ mice that could indicate an increased activation with improved cell survival via inhibition of apoptosis (Figure 2C, E, Suppl. fig. 2A, C).

Principal component analysis (PCA) of the DNA methylation data revealed clear separation of BMDM samples based on genotype (Dnmt3a^fl/fl^ vs Dnmt3a^ΔLysM^, Figure 3A, Suppl. fig. 3A). Moreover, samples from Dnmt3a^fl/fl^ mice clustered further according to age. While Dnmt3a^ΔLysM^ samples from young mice showed partial overlap, they remained distinct from aged Dnmt3a^fl/fl^ BMDMs, suggesting that Dnmt3a deficiency may elicit epigenetic changes comparable to, or exceeding, those induced by age-associated low-grade inflammation (inflammaging)^11,26^(Suppl. fig. 3A). Dnmt3a^ΔLysM^ samples from old mice, on the other hand, separated even further. According to polarization status, resting, M1 and M2 BMDMs for each genotype displayed an interspersed distribution among each genotype (Figure 3A). Overall, Dnmt3a loss induced hypomethylation on approximately 20,000 sites for all Dnmt3a^ΔLysM^ BMDM groups compared to their Dnmt3a^fl/fl^ counterparts (with almost half of them shared across polarization and age status), while hypermethylation was present in 9,500-15,000 sites (Figure 3B, Suppl. fig. 3B-C). In addition to sharing approximately 50% of hypomethylated sites across all samples (9,723 sites), resting

BMDMs from young mice exhibited the highest number of both unique hypo- and unique hypermethylated regions. (Figure 3B, Suppl. fig. 3B-C). When the differentially methylated positions (DMP) were classified, promoters and genes were more frequently hypomethylated than CpG islands (Figure 3C, Suppl. fig. 3D-E). Interestingly, promoter and genic regions in young M1 and aged M2 Dnmt3a^ΔLysM^ BMDMs exhibited a higher frequency of hypomethylation. Promoters were particularly affected in young M1 and M2 Dnmt3a^ΔLysM^ BMDMs and often showed shared methylation patterns across all groups. (Figure 3C, Suppl. fig. 3D-E).

To link the Dnmt3a-KO genotype to transcriptional dysregulation, we integrated RNA-seq results with those from DNA methylation. Correlation of differentially methylated positions (DMP, mostly hypomethylated) with differentially expressed genes (DEG) in cis^27^ highlighted two clusters of genes that increased expression in M2-polarized Dnmt3a^ΔLysM^ BMDMs, especially during ageing (Figure 3D-G, Figure 3H clusters 1 and 2). These genes relate to cell adhesion and migration ( *Itgb3, Itgb8, Cttn, Nectin1, Arhgef18, Myadm*), iron transport ( *Slc40a1*), macrophage stimulation and cytokine signaling ( *Mst1, Il7r, Palm3, Nlrc4*) and negative regulation of apoptosis ( *Uaca, Shc1*). Also increased in M2-polarized BMDMs from Dnmt3a^ΔLysM^ mice (mostly from young mice) are additional genes that increase expression during M1 polarization (independent of the genotype and age) which are related to cell proliferation and negative regulation of apoptosis (Figure 3H, cluster 3). Interestingly, genes governing inflammatory response (Nos2), Ca2^+^ signaling ( *Camk2d*), PI3K signaling ( *Nyap1*), and cell cycle ( *Mak, Lzts2*, and *Pkn3*) increase expression during M1 polarization in an age-dependent fashion in Dnmt3a^ΔLysM^ BMDMs (Figure 3H, cluster 4).

To uncover potential regulatory mechanisms, we inferred transcription factor binding sites (TFBS) associated with gene expression changes in BMDMs lacking Dnmt3a. These TFBS correspond to cell differentiation, including myeloid lineage differentiation, and MHC type II and cytokine/chemokine expression (Tcf12, Gata1, Ets1, and Ctcf, in addition to Tal1 in young M1/M2) (Figure 3I). Also, other transcriptions factors related to TGF-β signaling (p300, MyoD), binding to histone deacetylases HDAC (p300, Ctcf), circadian rhythm regulation (Bhlhe40) (Figure 3F). Of note, TFBS related to proto-oncogene Myc (Maz) and to hematopoiesis and/or cancer (Tal1, Ets1, Ctcf) could also be involved in the development of pathogenic conditions due to absence of *Dnmt3a* (Figure 3H-I). Many of these TFBSs were common for all polarization status and age, indicating genotype-specific alterations (Figure 3I).

Taken together, our results indicate *Dnmt3a* depletion induces BMDM reprogramming that affects cell morphology and adhesion, cell proliferation, and are associated with skewed M1 and M2 differentiation of BMDMs in vitro.

### M2 polarized macrophages lacking *Dnmt3a* expression are pro-phagocytic and hamper epithelial regeneration in vitro

Given that metabolic, cytoskeletal, and mitochondrial dysregulation may reflect impaired BMDM polarization, we assessed arginase (M2) and nitric oxide synthase (NOS; M1) activities, alongside analyses of cell morphology and phagocytic function. When assessing polarization, neither arginase nor NOS activities differed in cells from young mice (Figure 4A-B). However, M2-polarized macrophages from aged Dnmt3a^ΔLysM^ mice displayed diminished arginase (M2) activity compared to their aged WT counterparts (Figure 4A). Conversely, arginase activity was significantly elevated in M1 macrophages from aged Dnmt3a^ΔLysM^ mice (Figure 4A). Both M1 and M2 polarized macrophages from aged Dnmt3a^ΔLysM^ mice showed an increased NOS enzymatic activity compared to those from aged WT mice (Figure 4B). Resting Dnmt3a^fl/fl^ macrophages showed the slightly elongated shape typical of inactive cells, while resting Dnmt3a^ΔLysM^ macrophages had a higher proportion of rounded cells, resembling active M1 morphology (Figure 4C–D). M1 macrophages from both genotypes were predominantly rounded. In contrast, M2 macrophages from Dnmt3a^fl/fl^ mice displayed the expected elongated morphology, whereas over 50% of M2 macrophages from Dnmt3a^ΔLysM^ mice showed a disorganized structure and failed to fully adopt the M2-like shape. In vitro polarized BMDMs from aged mice behaved similarly to those from young mice (Figure 4C-D).

**Figure 4.**
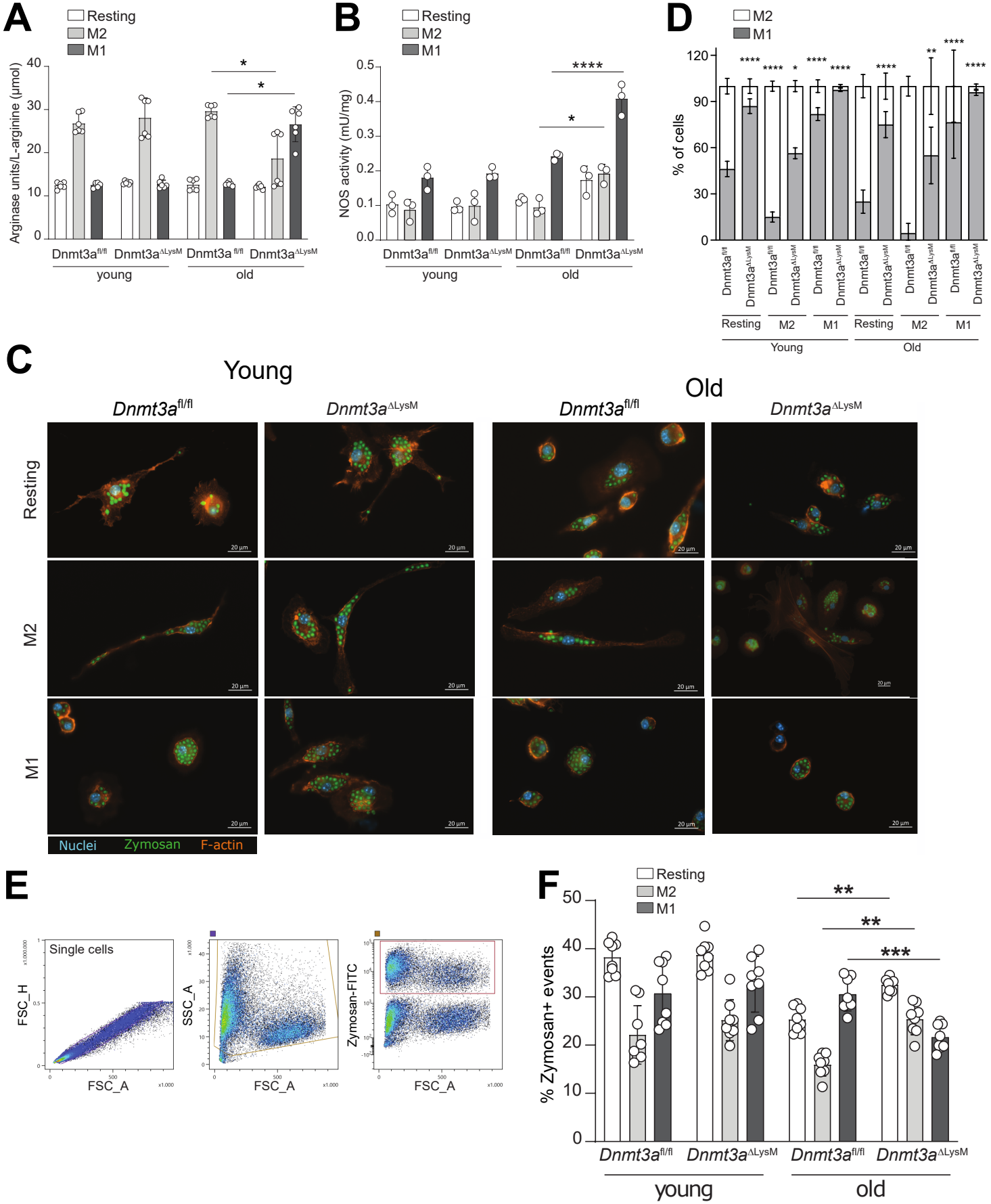
Depletion of Dnmt3a skews M2 BMDM polarization inducing an M1-like behavior in terms of morphology, enzymatic activity and phagocytosis. (A-B) Arginase (A) and nitric oxide synthase NOS (B) enzymatic activities on in vitro polarized BMDM from young (8-10 weeks old) and aged (>34 weeks old) Dnmt3a^fl/fl^ and Dnmt3a^ΔLysM^ mice. (C) Representative images of cytoskeletal disposition and zymosan-FITC engulfed particles by in vitro polarized BMDMs from young and old Dnmt3a^fl/fl^ and Dnmt3a^ΔLysM^ mice. Zymosan-FITC particles are shown in green, while F-actin cytoskeletal filaments stained with phalloidin are shown in orange and nuclei stained with DAPI are shown in blue. Scale bars represent 20μm. (D) Percentage of cells displaying M1 and M2 morphologies. Cells displaying a rounded, compact morphology were considered as M1, and cells displaying an elongated, spindle-like morphology were considered as M2. A minimum of 10 optic fields from 3 technical replicates were analyzed. (E) FACS gating strategy to exclude doublets and cell debris (left) and dead cells (center) to identify zymosan-FITC^+^ cells (right panel) from polarized BMDM. (F) Percentage of zymosan-FITC^+^ cells from polarized BMDM from young (8-10 weeks old) and old (>34 weeks old) Dnmt3a^fl/fl^ and Dnmt3a^ΔLysM^ mice assessed by FACS.

Further assessment of macrophage functionality by means of phagocytic activity using zymosan particles, showed that both Dnmt3a^fl/fl^ and Dnmt3a^ΔLysM^ young macrophages display a phagocytic behaviour according to their culture condition (resting macrophages phagocyte upon foreign particle encounter, M2 macrophages with limited activity, and M1 macrophages being prone to phagocytosis, Figure 4C-F). Macrophages from aged mice display an overall decreased phagocytotic capacity (Figure 4C-F), which aligns with previous findings in other *in vivo* mouse models^28^ and in human monocyte-derived dendritic cells *in vitro*^29^. However, resting and M2 macrophages from Dnmt3a^ΔLysM^ mice exhibited significantly enhanced phagocytic activity compared to their aged Dnmt3a^fl/fl^ counterparts, while phagocytosis decreased in M1 Dnmt3a^ΔLysM^ macrophages (Figure 4C-F). Together, the results suggest an age-dependent effect on morphological and phagocytic properties in Dnmt3a^ΔLysM^ macrophages towards a more active state.

Next, we assessed the regenerative capacity of macrophages by co-culturing scratch-wounded intestinal epithelial ModeK monolayers with polarized macrophages using transwell inserts. (Figure 5A). Within 36 h, ModeK cells almost completely closed the scratch lesion when cocultured with resting Dnmt3a^fl/fl^ macrophages from young mice. Co-cultures with resting Dnmt3a^ΔLysM^ macrophages displayed significantly reduced closure (Figure 5B-C). Comparing the effects of M1 and M2 co-cultures, we showed that M2 polarized Dnmt3a^fl/fl^ macrophages induced an almost complete wound closure while cells co-cultured with M2 Dnmt3a^ΔLysM^ macrophages failed to close the gap (Figure 5D-E). ModeK cells cocultured with M1 macrophages displayed the least defect closure, which even more pronounced in co-cultures with M1 Dnmt3a^ΔLysM^ macrophages (Figure 5D-E). Finally, we also monitored organoid-derived monolayer formation using transepithelial electrical resistance (TEER) measurement during coculture with polarized macrophages. While organoid monolayers reached up to 500 Ωcm^2^ after 12 days of coculture with M2 Dnmt3a^fl/fl^ macrophages, a maximum TEER value of 380 Ωcm^2^ was measured when exposed to M2 Dnmt3a^ΔLysM^ macrophages (Figure 4K). On the other hand, M1 macrophages prevented correct organoid-derived monolayer formation, an effect that was further aggravated with Dnmt3a^ΔLysM^ -derived macrophages (Figure 5F).

**Figure 5.**
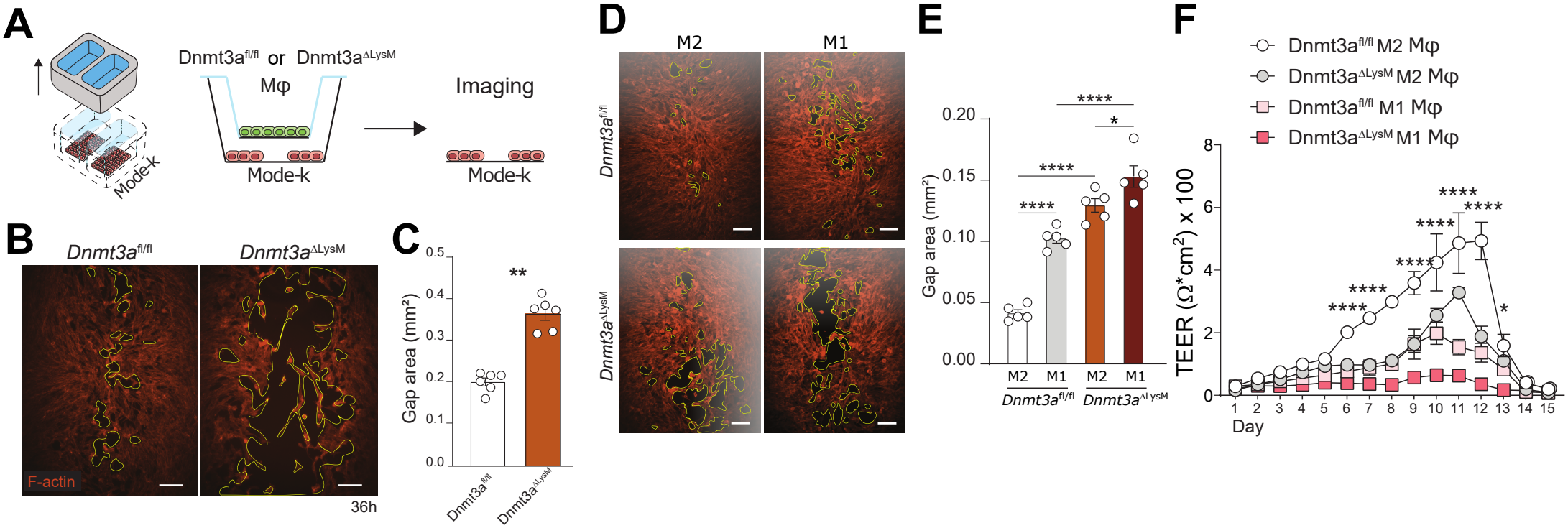
Dnmt3a-depleted M2 BMDMs prevent intestinal epithelial regeneration. (A) Schematic representation of the 36 h coculture of wounded intestinal epithelial ModeK monolayers with polarized macrophages in transwell inserts. (B) Representative images of ModeK monolayers exposed to resting BMDMs from Dnmt3a^fl/fl^ and Dnmt3a^ΔLysM^ mice for 36 h. F-actin cytoskeletal filaments stained with phalloidin are shown in orange. (C) Quantification of wound closure according to the area of the gap displayed in ModeK monolayers after coculture with resting BMDM from Dnmt3a^fl/fl^ and Dnmt3a^ΔLysM^ mice for 36 h. (D) Representative images of ModeK monolayers exposed to M1- and M2-polarized BMDMs from Dnmt3a^fl/fl^ and Dnmt3a^ΔLysM^ mice for 36 h. F-actin cytoskeletal filaments stained with phalloidin are shown in orange. (E) Quantification of wound closure according to the area of the gap displayed in ModeK monolayers after coculture with M1- and M2-polarized BMDM from Dnmt3a^fl/fl^ and Dnmt3a^ΔLysM^ mice for 36 h. (F) Transepithelial electrical resistance (TEER) measurements of organoid-derived intestinal epithelial monolayers cocultured with polarized BMDM from Dnmt3a^fl/fl^ and Dnmt3a^ΔLysM^ mice for 15 days. In all cases, BMDM were polarized using 20 ng/mL m-CSF for the resting program, 100 ng/mL LPS and 50 ng/mL IFN-g for the M1 program, and 50 ng/mL IL-4 and 50 ng/mL IL-13 for the M2 program. Zymosan-FITC particles were fed to polarized BMDM for 3 h in engulfment phagocytic experiments. Scale bars represent 20 μm in A and E, and 50 μm in F and H. All results are based on n = 5–8 biological replicates per group, except for panel K (n=3 per group). Data are represented as mean ± SEM. Statistically significant differences are shown as *p<0,05, **p<0,002, ***p<0,0002 ****, p<0,0001.

Altogether, M2 macrophages lacking *Dnmt3a* expression are more prone to display an activated rounded morphology characteristic of M1 macrophages and compromise wound healing properties. Also, with age, *Dnmt3a*-deleted M2 macrophages improve their phagocytic activity. In comparison, M1 macrophages seem to attenuate their phagocytosis with age while retaining proinflammatory effects that prevent adequate epithelial cell proliferation and wound closure.

### Mice lacking Dnmt3a expression in myeloid cells only show subtle alterations in experimental colitis, however spontaneously develop severe hepatosplenomegaly and signs of liver fibrosis with increasing age

Given the macrophage disrupted polarization and its role in epithelial homeostatic disruption, we explored the effects of macrophage *Dnmt3a* depletion during inflammation *in vivo*. 8-10-week-old Dnmt3a^fl/fl^ and Dnmt3a^ΔLysM^ female mice were exposed to 4% DSS for 5 days, allowing a recovery period of 7 days before sacrifice. Weight loss reached up to 19%, indicating efficient induction of colitis (Suppl. fig. 4A). Body weight loss was not significantly altered between Dnmt3a^fl/fl^ and Dnmt3a^ΔLysM^ mice; however, a slight increase in the disease activity index was evident in Dnmt3a^ΔLysM^ mice (Suppl. fig. 4A-B). All other physiological measures including caecum, spleen, liver or kidney weight, small intestine and colonic lengths and H&E histology score did not diverge between the experimental groups (Suppl. fig. 4C-I). Immune cell composition of neither spleen nor intestinal lamina propria differed (Suppl. fig. 4J-K).

To further assess the impact of prolonged inflammatory stress and involvement in colitis-associated carcinogenesis, we conducted a similar but longer experiment consisting of three cycles of DSS-recovery, combined with Azoxymethane (AOM) i.p. injections at the beginning and mid-term of the procedure. Dnmt3a^ΔLysM^ mice displayed an increased body weight at the experimental endpoint of 14 weeks (Suppl. fig. 5A-B). When tissues where examined, small intestinal and epididymal white adipose tissue (EWAT) weights of Dnmt3a^ΔLysM^ mice were increased, with histological evaluation revealing an increased colonic histological score compared to Dnmt3a^fl/fl^ mice (Suppl. fig. 5C-K). Immune cell composition in the spleen or the small intestinal and colonic lamina propria remained unchanged (Suppl. fig. 5L-N), suggesting Dnmt3a deficiency increases tissue susceptibility to chronic inflammation without altering overall immune cell proportions. Tumorigenesis was lacking, which could be due to the limited efficiency AOM could exert in C57BL/6J strain and in >6 week-old female mice, and the relatively low DSS dosage of 1-1.5%.^30^

Since *Dnmt3a*-dependent disruption in macrophage polarization seemed to be a cumulative, age-related event, we next sought to phenotype aged Dnmt3a^ΔLysM^ mice (>34 weeks-old). While young Dnmt3a^ΔLysM^ mice presented normal behaviour, offspring numbers, and organ morphology and anatomy; aged mice exhibited an increased abdominal enlargement and body weight resulting in impaired motility due to spontaneous hepatosplenomegaly (Figure 6A-G). Intestinal histology and intestinal stem cell proliferation remained normal in both young and aged Dnmt3a^ΔLysM^ mice (Figure 6H-I). However, liver and spleen histological architectures were severely altered in aged Dnmt3a^ΔLysM^ with marked expansion of lymphoid follicles and increased proliferation of mononuclear cells in the red and white pulp (Figure 6H-I). When splenic cells were investigated in detail, we found fewer HSCs in aged Dnmt3a^fl/fl^ control mice compared to young mice with the same genotype (Figure 7A-B). While HSC cell numbers remained normal in young Dnmt3a^ΔLysM^ mice, these were increased in aged Dnmt3a^ΔLysM^ mice and reached the same percentage of events as in young mice (Figure 7B). In addition, young Dnmt3a^ΔLysM^ mice displayed fewer macrophages compared to young Dnmt3a^fl/fl^ mice (Figure 7C). Age increased cell frequencies of T-cells and cytotoxic CD8^+^ T-cells, while decreasing CD11c^+^ dendritic cells (Figure 7D-G). In spleens from aged Dnmt3a^ΔLysM^ mice, T-cells, CD4^+^ helper T-cells and macrophages were decreased, whereas dendritic cells increased compared to aged Dnmt3a^fl/fl^ mice (Figure 7D-G). Neutrophils, natural killer, and B-cells remained comparable among all experimental groups (data not shown). On the other side, livers from Dnmt3a^ΔLysM^ mice were hardened due to vascularization with altered matrix organization according to Azan-based fibrotic score (figure 8) and modest mature collagen I deposition according to Ishak^31^ Sirius staining score (figure 8B-C). When liver gene expression was assessed, PCA from RNA-seq data separated samples according to sex and genotype, grouping Dnmt3a^fl/fl^ and Dnmt3a^ΔLysM^ male samples closer than their female counterparts (figure 8D). Of the 3402 DEGs, 2177 were upregulated and 1225 downregulated in Dnmt3a^ΔLysM^ mice (figure 8E). While upregulated genes related to proinflammation driven by IL-1β, IL-6, IFN-γ, and TNF, downregulated genes related to alterations in metabolism (fatty acid oxidation and biosynthesis, amino acid catabolism) and oxidative stress (mitochondrial respiratory chain assembly, glutathione metabolism) (figure 8F).

**Figure 6:**
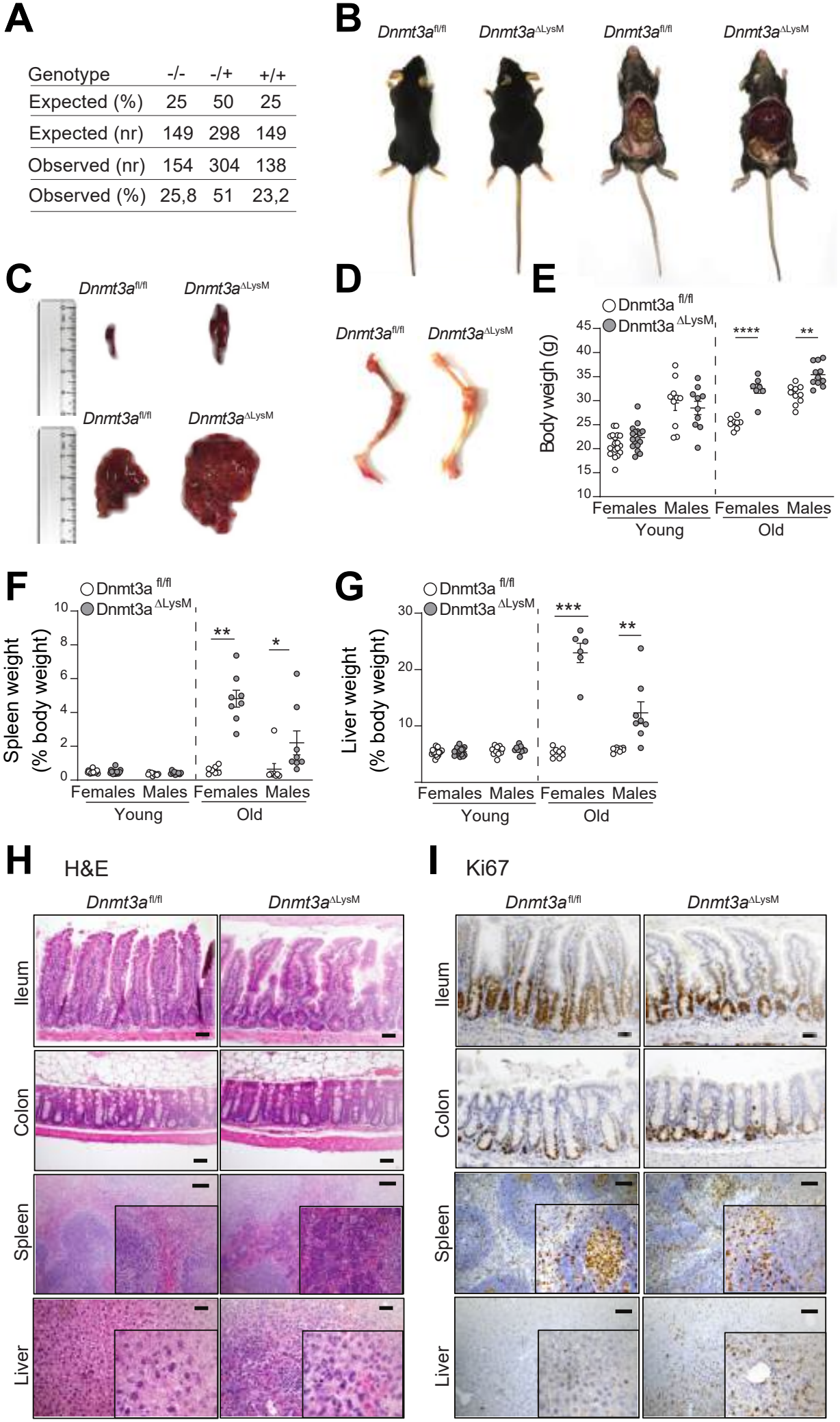
Aged Dnmt3a^ΔLysM^mice suffer from hepatosplenomegaly and display hyperproliferation of splenic and hepatic cells. (A) Birth rates of Dnmt3a^fl/fl^ and Dnmt3a^ΔLysM^ mice compared to expected mendelian ratios. (B-D) Representative images of body (B), spleen and liver (C) and femur and tibial (D) morphology of >34 weeks-old Dnmt3a^fl/fl^ (left) and Dnmt3a^ΔLysM^ (right) mice. (E-G) Weight of bodies (E), spleens (F), and livers (G) of young (8-12 weeks-old) and aged (>34 weeks-old) Dnmt3a^fl/fl^ and Dnmt3a^ΔLysM^ mice. (H-I) Representative images of haematoxylin and eosin staining (H) and proliferative marker Ki67 immunohistochemistry (I) of ileal, colonic, splenic, and hepatic tissue from >34 weeks-old Dnmt3a^fl/fl^ (left) and Dnmt3a^ΔLysM^ (right) mice. Scale bars represent 100 μm. Results are based on 10 male and 18 female 8–10-week-old mice, and 8 male and 5-6 female >34 week-old mice per genotype. Data are represented as mean ± SEM. Statistically significant differences are shown as *p<0,05, **p<0,002, ***p<0,0002 ****, p<0,0001.

**Figure 7:**
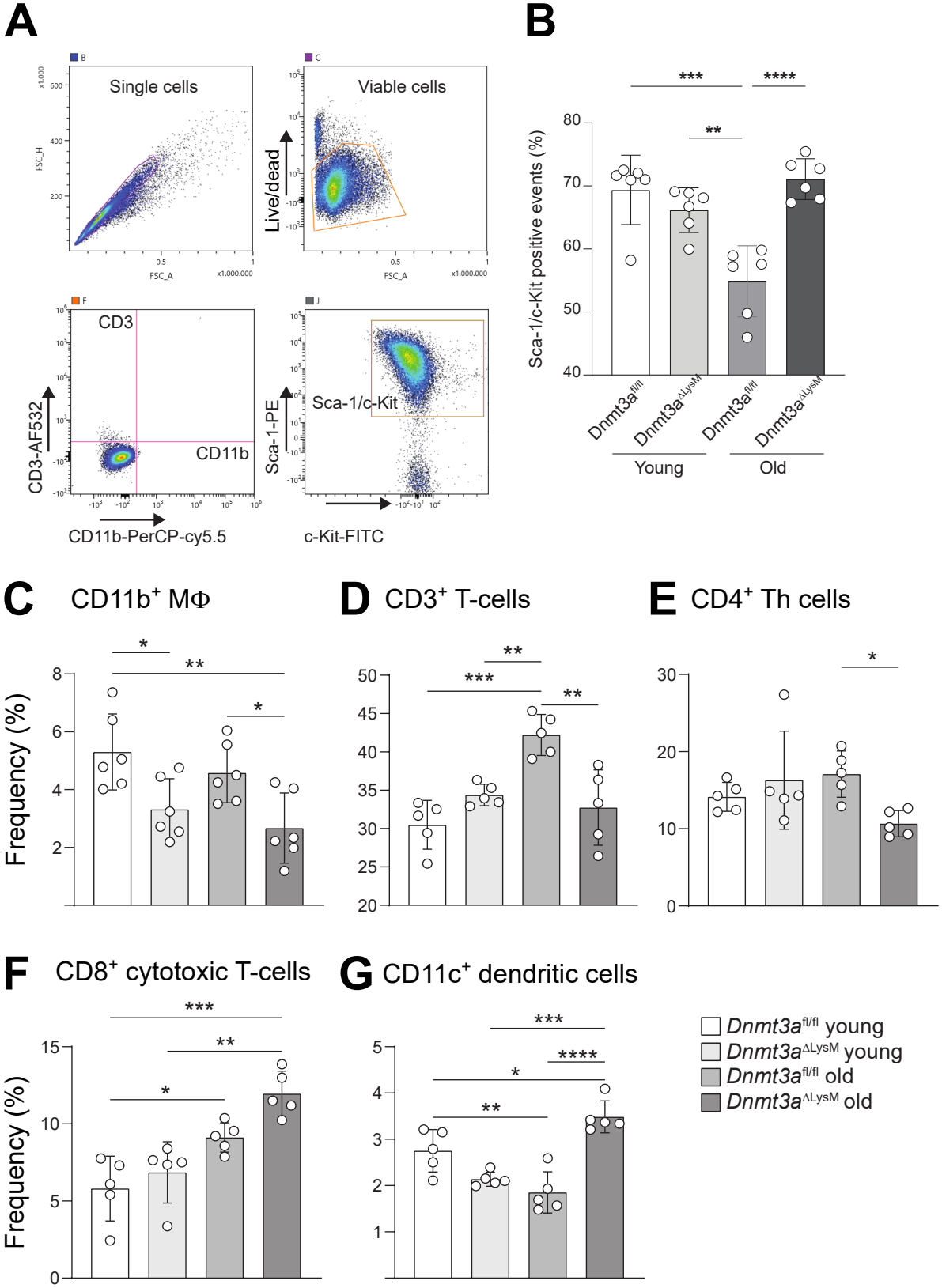
Lack of *Dnmt3a* expression in myeloid cells results in increased numbers of haematopoietic stem cells, cytotoxic T-cells, and dendritic cells in spleens from aged Dnmt3a^ΔLysM^mice. (A) FACS gating strategy to exclude doublets, cell debris and dead cells (upper panels) to identify hematopoietic stem cells (HSC: CD3^-^ CD11b^-^ Sca-1^+^ c-Kit^+^, lower panels) from MACS-isolated splenic HSC. (B) Percentage of splenic HSC from young (8-10 weeks old) and old (>34 weeks old) Dnmt3a^fl/fl^ and Dnmt3a^ΔLysM^ mice assessed by FACS. (C-G) Percentage of macrophages (CD11b^+^), T-cells (CD3^+^), T helper cells (CD4^+^), cytotoxic T-cells (CD8^+^), and dendritic cells (CD11c^+^) from spleens from young (8-10 weeks old) and old (>34 weeks old) Dnmt3a^fl/fl^ and Dnmt3a^ΔLysM^ mice assessed by FACS. All results are based on n = 5-6 mice per group. Data are represented as mean ± SEM. Statistically significant differences are shown as *p<0,05, **p<0,002, ***p<0,0002 ****, p<0,0001.

**Figure 8:**
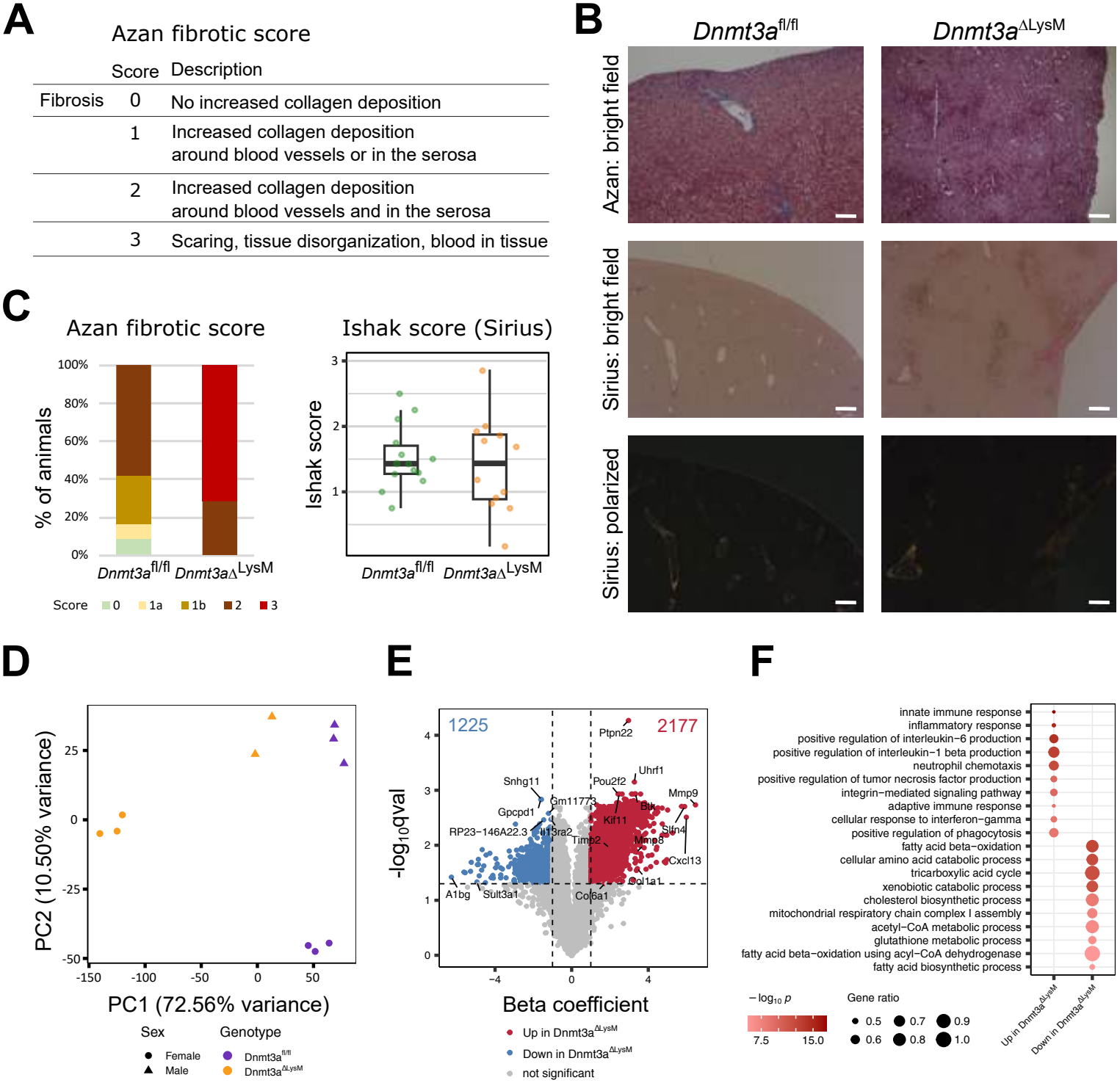
Livers from aged Dnmt3a^ΔLysM^mice display increased collagen deposition. (A) Criteria for histologic fibrotic scoring in liver based in Azan staining. (B) Representative images of matrix reorganization and fiber deposition (Azan staining) and mature collagen deposition (Sirius staining under bright field and polarized light microscopy) in livers from >34 weeks-old Dnmt3a^fl/fl^ (left) and Dnmt3a^ΔLysM^ (right) mice. (C) Fibrotic scores based on Azan (left) and Sirius (right) staining of livers according to Azan fibrotic score in A and Ishak *et al*.^31^ score system from >34 weeks old Dnmt3a^fl/fl^ and Dnmt3a^ΔLysM^ mice. Scale bars represent 50 μm. Results are based on 8 male and 6 female Dnmt3a^fl/fl^ and 7 male and 5 female Dnmt3a^ΔLysM^ mice of >34 week-old. Data are represented as mean ± SEM. (D) Principal component analysis (PCA) of liver transcriptome from aged (>34 weeks-old) Dnmt3a^fl/fl^ and Dnmt3a^ΔLysM^ mice. (E) Maaslin model volcano plot (accounting for sex and genotype) displaying upregulated differentially expressed genes (DEGs) in red and downregulated DEGs in blue for Dnmt3a^ΔLysM^ samples. (F) Gene ontology biological processes enriched in genes differentially expressed in livers from aged Dnmt3a^ΔLysM^ mice compared to Dnmt3a^fl/fl^ mice. Dot sizes indicate the ratio of significant DEGs to annotated genes in each term and colour corresponds to the p-value of enrichment. Analyses shown in D-F were conducted on liver samples from n=3 female Dnmt3a^fl/fl^, n=3 female Dnmt3a^DLysM^, n=3 male Dnmt3a^fl/fl^, and n=2 male Dnmt3a^DLysM^ mice; total n=11.

In summary, lack of *Dnmt3a* expression in myeloid cells surprisingly does not lead to significantly altered sensitivity for experimental DSS colitis in young mice. When Dnmt3a^ΔLysM^ mice age, splenic HSC propagation imbalances cell differentiation benefitting dendritic cell expansion over that of macrophages and CD4^+^ helper T-cells.

## DISCUSSION

The mechanisms by which reduced immune activation thresholds contribute to tolerance breakdown in inflammatory bowel disease (IBD) remain poorly defined. Our findings identify DNMT3A as a regulator of macrophage responses to polarizing cues. Dnmt3a-deficient BMDMs incompletely acquired IL-4/IL-13-associated features and provided reduced support for epithelial gap closure and barrier formation. These effects were accompanied by condition-dependent changes in phagocytosis and enzyme activity. In vivo, myeloid Dnmt3a deletion had limited effects on acute DSS colitis but was associated with greater histological injury in the AOM/DSS protocol and spontaneous hepatosplenomegaly in older mice.

Methylation events are, indeed, crucial for immune cell homeostasis. Monocyte differentiation into immature and then mature dendritic cells has been linked to dynamic changes in DNMT expression and methylation patterns.^32^ In T-cells, *DNMT3A* governs Th cell differentiation and directs early CD8^+^ T-cell effector and memory fate decisions while demethylation unstabilizes FoxP3 expression and disrupts immunoregulation.^20,33–35^ Similarly, B- and plasma cells lacking *Dnmt3a* and *Dnmt3b* are more active and proliferative upon immunization.^36^ Interestingly, *Dnmt3a* ablation prevents Ifng promoter methylation; hence, allowing IFN-γ production in Dnmt3a-deficient Th1, Th2, Th17 and even Treg cells.^20,37^ Here, we show that Dnmt3a-deficient bone marrow-derived macrophages (BMDMs) exhibit an age-dependent decrease in arginase activity and a concomitant increase in nitric oxide production, indicating a shift away from M2 polarization. Interestingly, previous studies have shown that mutations in the epigenetic regulators DNMT3A and TET2 can paradoxically enhance ARG1 expression and arginase activity, features typically associated with M2, not M1, macrophages.^38^

We found that phagocytic capability of Dnmt3a-depleted resting and M2 polarized BMDM from aged mice increased, which is in agreement with that from human monocytes and monocyte-derived M2-polarized macrophages after exposure to hypomethylating agent decitabine.^39^ *Dnmt3a* also methylates loci of the gene coding for phagosome cargo sequester LC3 ( *MAP1LC3*), which is involved in pathogen LC3-associated phagocytosis (LAP) and autophagy.^40,41^ Thus, Dnmt3a seems key not only for pathogen engulfment and immune activation, but also for autophagy, a key cellular process involved in IBD pathogenesis and disease progression.^42^ In another study, diabetic wound macrophages were shown to have destabilized *DNMT3a/b* expression which was associated with increased COX-2 and prostaglandin E2 levels. The authors postulated that this could contribute to increased cytokine production and perturbed wound healing.^43^ Indeed, we can show that deletion of Dnmt3a in M2-polarized BMDM impairs their beneficial effect on wound healing of intestinal epithelial monolayers *in vitro* scratch assays, confirming the prior hypothesis. Not only macrophage wound healing is affected, but evidence also points to a role of methylation and DNMTs in neutrophil-mediated early wound healing disruption and neutrophil extracellular trap (NET) release.^44,45^

Mice lacking myeloid *Dnmt3a* expression spontaneously developed hepatosplenomegaly and initiated liver fibrosis with age as well as dysbalanced immune cell differentiation. *Dnmt3a* has been found to be essential for hematopoietic stem cell differentiation^46^, but it seems it can also maintain HSC hyperproliferation.^23,47^ Also for adult mesenchymal stem cells (MSC), demethylation improves proliferation while decreasing oxidative stress.^48^ Interestingly, *DNMT3A* also impacts hepatic stellate cell proliferation, activation and autophagy, and associates with liver fibrosis.^49^ While *DNMT3A* has been recognized as a risk gene for IBD and IBD patients risk for metabolic-associated fatty liver disease and liver fibrosis,^14,50^ it remains to be elucidated whether *DNMT3A* could promote – or at least initiate – liver fibrosis in IBD. Supporting this hypothesis, livers from mice lacking Dnmt3a myeloid expression show signs of matrix reorganization and initial fibrotic activity, including transcriptional alterations of key liver metabolic pathways, oxidative stress and inflammation.

In summary, our results confirm that *Dnmt3a* is required for physiological macrophage polarization as its depletion leads to a pro-inflammatory skewing, especially during M2-polarization with IL-4 and IL-13. Myeloid lack of *Dnmt3a* induces spontaneous development of hepatosplenomegaly, which is led by HSC hyperproliferation. Further profiling macrophagic IBD epigenetic fingerprint could provide novel disease-progression biomarkers and highlight new druggable targets.

## METHODS

### Mice and induction of experimental colitis

Dnmt3a^fl/fl^ mice of C57BL/6J background were generated by the European Mouse Mutant Archive EMMA (Infrafrontier, München, Germany). Briefly, exons 13-17 were flanked by LoxP sites to enable its excision by Cre recombinase. Upstream exon 13, a distal loxP site was introduced with an FRT flanked neomycin selection cassette. The resultant mouse line was bred with Flp deleter-mice (C57BL/6-Tg(CAG-Flpe)2Arte, Taconic, Rensselaer, NY, USA) constitutively expressing Flp recombinase to remove the neomycin selection cassette, creating a Dnmt3a^fl/fl^ mouse in which Dnmt3a exons 13–17 were flanked by two loxP sites. These mice were crossed with mice containing a nuclear localized-Cre under the control of endogenous Lysozyme 2 ( *Lyz2*) promoter/enhancer elements that abolishes *Lyz2* gene function in myeloid cell lineage (LysM-Cre), resulting in LysM-cre^+/-^;Dnmt3a^fl/fl^ mice with myeloid-cell-specific Dnmt3a deletion (termed Dnmt3a^ΔLysM^ hereafter). For the baseline investigations male and female mice (age 8–12 and >34 weeks) were employed at a 1:1 ratio, distributed equally between Dnmt3a^fl/fl^ control and Dnmt3a^ΔLysM^ knockout groups. Experimental colitis was induced by administrating 4% dextran sodium sulfate (DSS, MP Biomedicals, Irvine, CA, USA) in autoclaved drinking water during a period of 5 days, followed by a recovery period of 7 days with clean water in 10-12-week-old female mice (8 mice/genotype). Chronic inflammation and carcinogenesis were induced by administrating three DSS cycles (1% for cycle 1, and 1.5% for cycles 2 and 3) with 2-week recovery periods between cycles, combined with 10mg/kg Azoxymethane (AOM) intraperitoneal (i.p.) injections at the beginning (week 1) and mid-term (week 7) of the procedure. Disease activity index (DAI) was used to monitor the severity of the treatment daily during experimental colitis. DAI is based on a 0 to 5 score on body weight loss (0, 1-4.99 %, 5-9.99 %, 10.19-99 %, or ≥ 20 %), stool consistency (formed, soft, very soft, unformed, or liquid), and presence of blood in stools (negative, slightly positive, positive, visible blood, or rectal bleeding).^51^

All mice were maintained in a specific pathogen-free facility with a 12⍰h light-dark cycle at 21 ± 2 °C and 60 ± 5 % humidity, and provided with food and water (or DSS) ad *libitum*. Mice were euthanized by cervical dislocation prior to sampling organs for histological and molecular analyses. Animal experiments were approved by the committee for animal welfare of the state of Schleswig-Holstein (V242-7224.121-33; V242-28372/2022 (29-4/22)).

### Intestinal lamina propria and epithelial cell dissociation

Small intestines and colons were harvested and flushed with PBS. Afterwards, they were cut longitudinally and rolled up in Swiss roll disposition for subsequent histological analyses or further cut into 0.5 cm pieces for organoid generation (see below) or cell isolation using the Lamina Propria Dissociation Kit for mouse (Miltenyi BioTech, Gaithersburg, MD, USA) following the manufacturer’s protocol. After dissociation and cleaning with a 30 μm cell strainer, CD11b^+^ cells were isolated using magnetic activated cell sorting (MACS) with CD11b microbeads and MS columns (Miltenyi Biotech). Cell populations were analyzed using spectral cytometry.

### Splenocyte and hematopoietic stem cell isolation

Spleens were collected in RPMI 1640 medium and mashed through a 40 μm cell strainer to isolate splenocytes. Splenocytes were obtained from the gradient interphase after adding 1.099 g/mL Ficoll solution (Sigma-Aldrich, St. Louis, MA, USA) and centrifuging at room temperature, RT (700rpm, 20min). Afterwards, splenocytes were washed twice in RPMI medium supplemented with 10 % fetal calf serum (FCS). Hematopoietic stem cells (HSC) were enriched using Lineage Cell Depletion Kit for mice (Miltenyi Biotech, Gaithersburg, MA, USA) following the manufacturer’s recommendations. Eluted HSC cell purity was verified using spectral cytometry.

### Bone marrow-derived macrophage (BMDM) harvest and macrophage differentiation and polarization

Bone marrow-derived macrophages (BMDM) were obtained from murine femurs and tibias of rear legs after removal of muscular and tendinous tissue and kept on ice cold RMPI 1640 medium (Invitrogen, Waltham, MA, USA). Apophyseal bone parts were cut to flush the bone marrow using a 26 G needle with BMDM culture medium (1:1 DMEM and macrophage serum-free medium supplemented with 10 % FCS, 5 % penicillin/streptomycin and 5 % amphotericin B) and filtered using a 70 μm cell strainer. BMDMs were differentiated in BMDM medium with 20 ng/mL macrophage colony-stimulating factor (m-CSF) for 5 days at 37 ºC and 5 % CO2. Adherent differentiated macrophages were washed and covered with ice-cold PBS to allow cell detachment for 5 min. Then, warm BMDM medium was added, and cells were scraped. After collection and centrifugation, cells were counted and seeded using BMDM medium with 50 ng/mL LPS and 50 ng/mL IFN-γ for M1 polarization, 50 ng/mL IL-4 and 50 ng/mL IL-13 for M2 polarization, or 20 ng/mL m-CSF for resting, unpolarized control conditions.

### Phagocytosis activity assessment

Macrophage phagocytosis activity was monitored for 3 h using Phagocytosis assay kit containing green zymosan particles (Abcam) following the manufacturer’s recommendations. Complementarily, cells were directly analyzed using spectral cytometry on a SA3800 spectral analyzer (Sony Biotech) or fixed with 4 % paraformaldehyde (PFA) and stained with phalloidin rhodamine and 41,6-diamidino-2-phenylindole (DAPI) to visualize F-actin filaments and cell nuclei, respectively, in a Leica laser SP5 scanning confocal microscope (Leica Microsystems).

### BMDM-induced wound healing assays with intestinal epithelial ModeK cells

ModeK cells were maintained in minimal essential medium (MEM) supplemented with 20 % FCS and passed using 0.25 % trypsin-ethylenediaminetetraacetic acid (EDTA) solution (Invitrogen). A total of 3 x 10^4^ cells were seeded in a 2-well culture insert (Ibidi, Gräfelfing, Germany) until cells reached confluency. Then, inserts were removed leaving a 500 μm gap between the cell monolayer from each chamber. ModeK medium was then refreshed and transwell inserts (Corning, Corning, NY, USA) containing previously polarized BMDMs were placed on top. ModeK proliferation and gap closure was monitored for 36 h using a Zeiss Axio Vert.A1 inverted microscope (Zeiss, Jena, Germany) and images were analyzed using ImageJ software (NIH, USA).

### Organoid culture and BMDM-induced epithelial barrier generation

Crypts Cwere isolated after colonic 0.5 cm pieces were incubateded with cold Ca^2+^/Mg^2+^-lacking 2.5mM EDTA solution for 40 min and mechanically dissociateded as described by Sato *et al*.^52^Crypts were recovered after centrifugation at 300 g and 4 ºC for 10 min, and filtering through a 100 μm cell strainer. Then, approximately 10 crypts per μL Matrigel® (Becton Dickinson, Franklin Lakes, NJ, USA) were left to polymerize for culture in organoid medium (50% L-WNR conditioned medium, with 50% 2x Advanced DMEM/F12 medium supplemented with 1 M HEPES, 1% Glutamax, 1% penicillin/streptomycin, 500 mM N-acetyl-Cysteine and 30% FCS). Organoids were recovered with ice cold PBS and reduced to single cell suspensions using TrypLE Express (Gibco) at 37 ºC for 2-3 min and filtered through a 70 μm cell strainer. A total of 1.5 x 10^5^ cells were seeded per transwell insert (Corning) using organoid medium. Epithelial cell-containing transwell inserts were then placed on top of polarized macrophage cultures with a cell density of 3 x 10^4^ cells/well in 24-well plates and monitored using a Millicell® ERS-2 Voltohmmeter (Merck, Darmstadt, Germany) for 15 days.

### Spectral cytometry

Up to 2 x 10^5^ cells were labelled Zombie Red Fixable Viability Kit (BioLegend, 1:1000) and non-specific Fc receptor blocked with anti-CD16/32 (BioLegend) at 4 ºC for 20 min and protected from light. After washing in PBS containing 1% FCS, antibodies were incubated at 4 ºC for 40 min protected from light.

Intestinal lamina propria, mesenteric lymph nodes (MLN) and splenic cells were used to detect immune cell subsets with antibodies against CD45 (1:100, FITC), CD11b (1:50, PerCP/Cy5.5), Lys6G (1:100, Brilliant Violet BV421), CD4 (1:100, BV711), CD19 (1:100, BV650), CD8 (1:100, BV786), CD11c (1:80, Pacific blue) (all from Biolegend, San Diego, CA, USA) and NK1.1 (1:50, PE) and CD3ε (1:100, Alexa Fluor 532) from eBioscience (ThermoFisher Scientific). In addition, splenic cells were used to verify HSC purity with antibodies against c-kit (1:60, FITC), Sca-1 (1:50, PE), and CD3ε (1:100, Alexa Fluor 532) antibodies from eBioscience, and CD11b (1:100, PerCP/Cy5.5), and CD19 (1:100, BV650) antibodies from Biolegend. Samples were analyzed in a SA3800 spectral cell analyzer and software v2.0.5 (Sony Biotechnology, Tokyo, Japan)

### Histology, immunohistochemistry, and immunofluorescence

Freshly harvested tissue was fixed in 10% w/v formalin at 4 ºC for 24 h. Then, tissues were dehydrated in increasing concentrations of ethanol until reaching 100%, and subsequently replaced with xylol substitute (Carl Roth, Karlsruhe, Germany) before paraffin embedding. Sections of 4⍰µm were deparaffinized with Xylol substitute and rehydrated in a decreasing ethanol series. Samples for hematoxylin and eosin (H&E) staining were then stained for 2-5min in hematoxylin and counterstained with 1% v/v eosin solution for 2min. Samples for apoptosis determination were subjected to Apop Tag Plus Peroxidase In Situ Apoptosis Detection kit (Merck Millipore, Darmstadt, Germany) according to the manufacturer’s protocol. Samples used to detect collagen and reticular connective tissue were stained with Azan staining kit (Morphisto, Offenbach am Main, Germany) following the manufacturer’s instructions. For immunohistochemistry (IHC), antigen retrieval was performed using boiling citrate buffer at pH 6.0 for 20 min, blocking of endogenous peroxidase using hydrogen peroxidase for 10 min, and blocking of antigens using 1 % bovine serum albumin (BSA) for 30 min. Primary anti-bromodeoxyuridin, BrdU (1:10 dilution, BD Pharmingen, Heidelberg, Germany) and anti-DNMT3A (1:100 dilution, Cell Signaling Technologies, Cambridge, UK) were incubated at 4 ºC overnight. IHC sections were washed and incubated with secondary antibody goat anti-mouse immunoglobulin G (1:1000; Jackson ImmunoResearch, Ely, UK) and 3,3’-diaminobenzidine (DAB) substrate with Vectastain ABC Kit (Vectorlab, Peterborough, UK). All slides were dehydrated and mounted using xylene-based mounting media. Images were acquired in a Zeiss Axio Imager Z1 microscope with ZEN 3.4 Blue edition software (Zeiss). Histological scores ranging 0-3 (absent, mild, moderate, or severe) were obtained based on mononuclear cell infiltration, crypt hyperplasia, epithelial injury/erosion, and 0-4 for transmural inflammation (absent, submucosal, one focus extending into muscularis and serosa, up to 5 foci extending into muscularis and serosa, or diffuse). Liver damage was scored using Azan staining ranging 0-3 (normal, increased collagen deposition on liver edge/serosa and/or around blood vessels, scaring and disorganization).

*Ex vivo* generated BMDMs were cultivated in 8-well chamber slides (1.0-1.5 x 10^4^ cells/well). Cells were washed, fixed in 4 % PFA in PBS, and permeabilized in 0.5 % Triton X-100 in PBS for 3-5 min. The unspecific binding sites were blocked using 5 % BSA for 30 min prior to primary antibody incubation overnight at 4⍰°C and secondary antibody for 2⍰h. DAPI (1:40,000) and DRAQ5 (1:5,000) were used to label cell nuclei while phalloidin rhodamine (1:500) was used to visualize F-actin filaments. Confocal images were acquired in a Leica laser SP5 scanning confocal microscope (Leica Microsystems, Wetzlar, Germany) and processed using ImageJ software.

### RNA isolation and mRNA sequencing (RNA-seq) analyses

Cells from *in vitro* assays were directly lysed on 350μL RLT buffer supplemented with 10% beta-mercaptoethanol. Tissues were snap frozen and kept at -80 ºC until RNA isolation. RNA was isolated with RNeasy Mini Kit including DNAse digestion using the RNAse-free DNAse set (all from Qiagen, Hilden, Germany) following the manufacturer’s recommendations for cells or tissue, respectively. Sequencing libraries were prepared according to the Illumina stranded messenger RNA (mRNA) sequencing protocol and sequenced (2 x 150 bp reads, median depth of 50 (37) million read pairs per BMDM (liver) sample) on an Illumina NovaSeq 6000 (all from Illumina, San Diego, CA).

The nf-core rnaseq pipeline version 3.3 (https://github.com/nf-core/rnaseq)^53^ was used to trim adapter and low-quality bases using Trim Galore v0.6.6 (http://www.bioinformatics.babraham.ac.uk/projects/trim_galore/). Trimmed reads were filtered for <20bp reads and mapped to the mouse genome GRCm38, gencode version M6 using STAR aligner v2.6.1d^54^. Reads from two young Dnmt3a^ΔLysM^ BMDM samples were excluded due to alignment to *Lyz2* and *Dnmt3a* exons 13-17 and data from one Dnmt3a^ΔLysM^ liver sample was removed due to read alignment to *Lyz2*. Quantification of transcript expression was performed using salmon v1.4.0^55^, and imported to gene-level counts with tximport v1.28.0^56^. Only genes with expression counts ≥5 in all BMDM samples of at least one combination of genotype, polarization status and age group (14,478 genes) were included in subsequent analyses. For liver samples, genes with expression counts ≥5 in all samples of at least one combination of genotype and sex (14,884 genes) were used in downstream analyses. In both BMDM and liver sample analyses, expression of genes that passed the respective filtering step were normalized across samples using DESeq2’s median of ratios method (v1.40.2^57^). Principal component analysis (PCA) was performed on variance stabilized expression counts.

Differentially expressed genes (DEG) between Dnmt3a^fl/fl^ and Dnmt3a^ΔLysM^ samples were obtained using Maaslin2 v1.14.1 linear mixed models^58^. For BMDM samples young and old mice were analyzed separately using polarization status and genotype as fixed effects and variance stabilized expression counts as response variable (significance by Benjamini-Hochberg false discovery rate (FDR) adjusted p-values <0.05). For liver samples sex and genotype were used as fixed effects (significance by Benjamini-Hochberg false discovery rate (FDR) adjusted p-values <0.05 and |beta coefficient| > 1). DEG expression heatmaps were created using ComplexHeatmap v2.16.0^59^. DEGs from BMDM samples were clustered through k-means clustering (repeated 100 times), and manually characterized based on gene ontology enrichment results.

### DNA isolation and methylation analyses

Differentiated BMDMs were homogenized in PBS supplemented with 1% proteinase K. DNA from lysates was obtained using DNeasy Blood and Tissue kit (Qiagen) according to the manufacturer’s instructions. Bisulfite conversion and methylation profiling were performed using Infinium© Mouse Methylation BeadChip, and analysed using the Bioconductor package RnBeads v2.20.0.^60^ Sites that overlapped with single nucleotide polymorphisms or had unreliable measurements were filtered out. Context-specific probes, probes on the sex chromosomes, and probes with missing values were also removed. In total, 14,351 out of 261,600 probes were filtered. The signal intensity values were normalized using the dasen method. PCA was performed on methylation intensity (β values) of all sites. Differentially methylated positions (DMPs), promotors, genes and CpG islands between Dnmt3a^fl/fl^ and Dnmt3a^ΔLysM^ samples were identified using the automatically selected rank cutoff of RnBeads. Transcription factor binding sites (TFBS) enriched in DMPs were identified using LOLA v1.30.0^61^.

### Transcriptome and DNA methylation integration analyses

For the integrated analysis of gene expression with DNA methylation, DMPs located 5 Kbp up- and downstream of transcription start sites of DEGs were firstly identified. Spearman’s rank correlation coefficient between each DEG normalized expression and DMP methylation intensity β values was calculated. A permutation approach was used to calculate statistical significance of correlations. TFBS enriched in significantly correlated DMP-DEG pairs were identified using LOLA v1.30.0^61^.

### Functional enrichment analysis

All gene ontology (GO) enrichment analyses were conducted using topGO v2.46.0, with genes with similar expression level as the universe set (Fisher.elim p-value calculated using the weight algorithm <0.05 was taken as significant)^62^.

### Reverse transcription quantitative polymerase chain reaction (RT-qPCR)

RNA was quantified using NanoDrop ND-2000 and reverse-transcribed with Maxima H minus First Strans cDNA synthesis kit (ThermoFisher Scientific, Waltham, MA, USA). TaqMan Gene Expression Master Mix and TaqMan® probes (Applied Biosystems, Darmstadt, Germany) were used for RT-qPCR in a VIIA 7 PCR system (ThermoFisher Scientific). Gene expression was obtained after relative -ΔCt quantification with reference gene Actb or Gadph. Each sample was analyzed in duplicate.

### Western blot

Proteins from cultured cells were obtained with RIPA buffer supplemented with 1% Halt Preotease and Phosphatase Inhibitor Cocktail following the manufacturer’s instructions (ThermoFischer Scientific), and quantified using DC protein assay (BioRad, Munich, Germany). Electrophoresis was performed in 7.5-12% acrylamide/bis-acrylamide gels and proteins were transferred to a polyvinylidene difluoride (PVDF) membrane in semi-dry Trans-Blot Turbo TM Transfer System (all from BioRad). Blockage was performed with 5% BSA or 5% non-fat dry milk in 0.1% Tween20 Tris-buffered saline solution (all from ThermoFisher Scientific). Primary antibodies against Dnmt3a (1:1000 in 5% non-fatty dry milk, overnight at 4 ºC, R&D Biosystems), arginine-1 (1:2000 in 5% BSA, overnight at 4 ºC, Invitrogen), inducible nitric oxidase iNos (1:1000 in 5% BSA, 1 h at RT, Abcam), and Gadph (1:800 in 5% non-fatty dry milk, 45 min at RT, Abcam) were incubated in blocking solution. Horseradish peroxidase (HRP)-labelled secondary antibodies against rabbit or mouse immunoglobulins were incubated for 1 h at RT. Chemiluminescent signal was visualized after 5min incubation with ECL substrate (GE Heatlhcare, Freiburg, Germany) in a ChemiDoc MP Imaging System with ImageLab v-6.1.0 (BioRad).

### Arginase activity assay

One million BMDMs were differentiated in 6-well plates for 4 days after differentiation to macrophages. After collection with ice cold PBS, scrapping and centrifugation as described above, cells were lysed in 100 μL 10 mM Tris-HCl buffer at pH 7.4 containing 1:100 pepstatin A, 1:100 Leupeptin, and 0.4% Triton X100 solution from the Arginase Activity Assay Kit (Sigma-Aldrich). Cell content was collected after centrifugation at 13,000 g for 10 min and arginase activity was quantified using Arginase Activity Assay Kit following the manufacturer’s instructions. Absorbance at 430 nm was read in an Infinite F200 pro plate reader (Tecan, Männedorf, Switzerland).

### Nitric oxide synthase assay

Five million differentiated BMDMs were homogenized in 200 μL cold nitric oxide synthase (NOS) Assay buffer. Cell content was obtained after 10 min centrifugation at 10,000 g and 4 ºC. Proteins were quantified using DC protein assay (BioRad). Afterwards, 200 μg proteins were diluted in NOS assay buffer and NOS activity was quantified using Nitric Oxide Synthase Activity Assay Kit (Abcam, Waltham, MA, USA).

### Statistical analyses

Non-parametric Mann Whitney test was used to analyze data. Data shown display mean ± standard error of the mean (SEM). A p-value of ≤ 0.05 was considered as statistically significant. Statistical analyses were performed and plotted using GraphPad Prism 9 (GraphPad Softwre Inc., La Jolla, CA, USA). Analyses of RNA sequencing and methylation data were performed using R v4.3.1. FDR correction for multiple testing was performed when applicable.

## Data availability

The raw and processed RNA-sequencing data and Infinium© Mouse Methylation BeadChip array data of BMDMs have been deposited in the NCBI GEO database under the accession numbers GSE286366 and GSE286367, respectively. RNA-sequencing data of liver samples have been deposited under the accession number GSE334527. All datasets can be accessed through the superseries GSE334528 and will be available upon peer-review and publication in a journal. Remaining data are available from the corresponding author on reasonable request.

## Code availability

All custom codes generated for RNA-seq, methylation and integration bioinformatics analyses can be found in https://github.com/Systems-Immunology-IKMB/DNMT3A_LysM_BMDM and will be available upon peer-review and publication in a journal.

## ACKNOWLEDGEMENTS

We thank Stefanie Rentzow, Tanja Klostermeier, and Maren Reffelmann for support with sample collection and IHC; Sabine Kock, Vivian Wegener and Dorina Oelsner for support with PCR and sequencing sample preparation; and Meike Hansen, Ronja Möhring and Sophie Reiher for support with mouse genotyping. Sequencing was performed at the Competence Centre for Genomic Analysis (CCGA) Kiel, which is funded by the German Research Foundation, DFG (Deutsche Forschungsgemeinschaft) and the state of Schleswig-Holstein.

## SUPPLEMENTARY FIGURE LEGENDS

**Supplementary figure 1.**
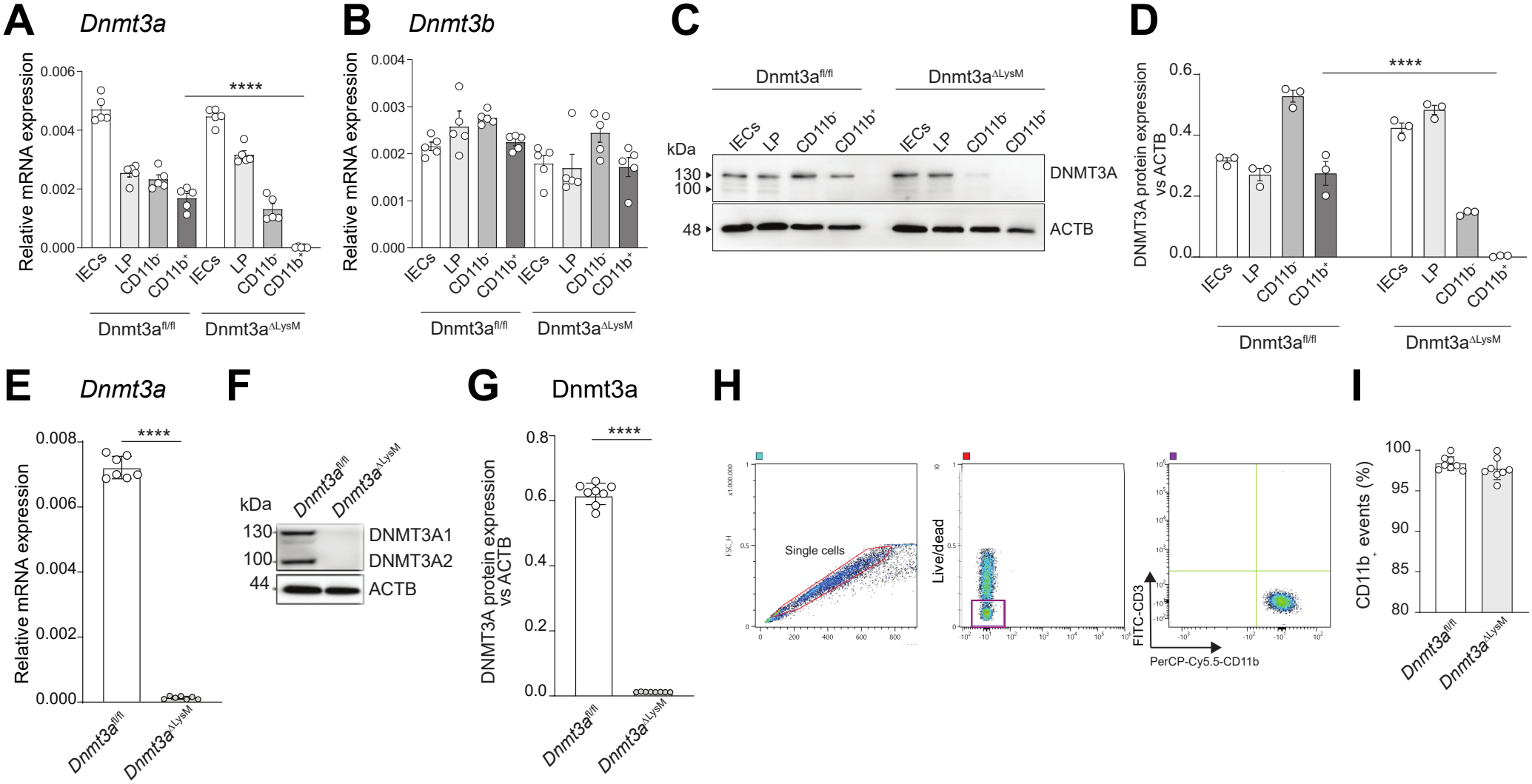
Deletion of Dnmt3a in myeloid cells from Dnmt3a^ΔLysM^mice shows no changes in cell proportions of CD11b+ events in *ex vivo* differentiated BMDM. (A-B) Relative mRNA expression levels of *Dnmt3a* (A) and *Dnmt3b* (B) in intestinal epithelial cells (IEC), lamina propria (LP), and MACS-isolated CD11b^-^ and CD11^+^ LP fractions of small intestines from Dnmt3a^fl/fl^ and Dnmt3a^ΔLysM^ mice. *Actb* was used as housekeeping gene for normalization. (C-D) Dnmt3a protein detection by western blot (C) and its relative semi-quantification (D) from 15mg of total protein from IEC, LP, and MACS-isolated CD11b^-^ and CD11^+^ LP fractions of small intestines from Dnmt3a^fl/fl^ and Dnmt3a^ΔLysM^ mice. Actb was used as reference control. (E) Relative mRNA expression levels of Dnmt3a from in vitro differentiated BMDM from Dnmt3a^fl/fl^ and Dnmt3a^ΔLysM^ mice. Actb was used as housekeeping gene for normalization. (F-G) Dnmt3a protein detection by western blot (F) and its relative semi-quantification (G) from 15mg of total protein from *in vitro* differentiated BMDM from Dnmt3a^fl/fl^ and Dnmt3a^ΔLysM^ mice (5-day exposure to 20 ng/mL m-CSF). Actb was used as reference control. (H) FACS gating strategy to exclude doublets and cell debris (left) and dead cells (center) to identify CD11^+^ cells (right panel) from differentiated BMDM. (I) Percentage of CD11+ cells from *in vitro* differentiated BMDM from Dnmt3a^fl/fl^ and Dnmt3a^ΔLysM^ mice assessed by FACS. All results are based on n = 5–8 animals per group. Dnmt3a^fl/fl^ samples are shown in white, Dnmt3a^ΔLysM^ samples are shown in grey. Data are represented as mean ± SEM. Statistically significant differences are shown as ****p<0,0001.

**Supplementary figure 2.**
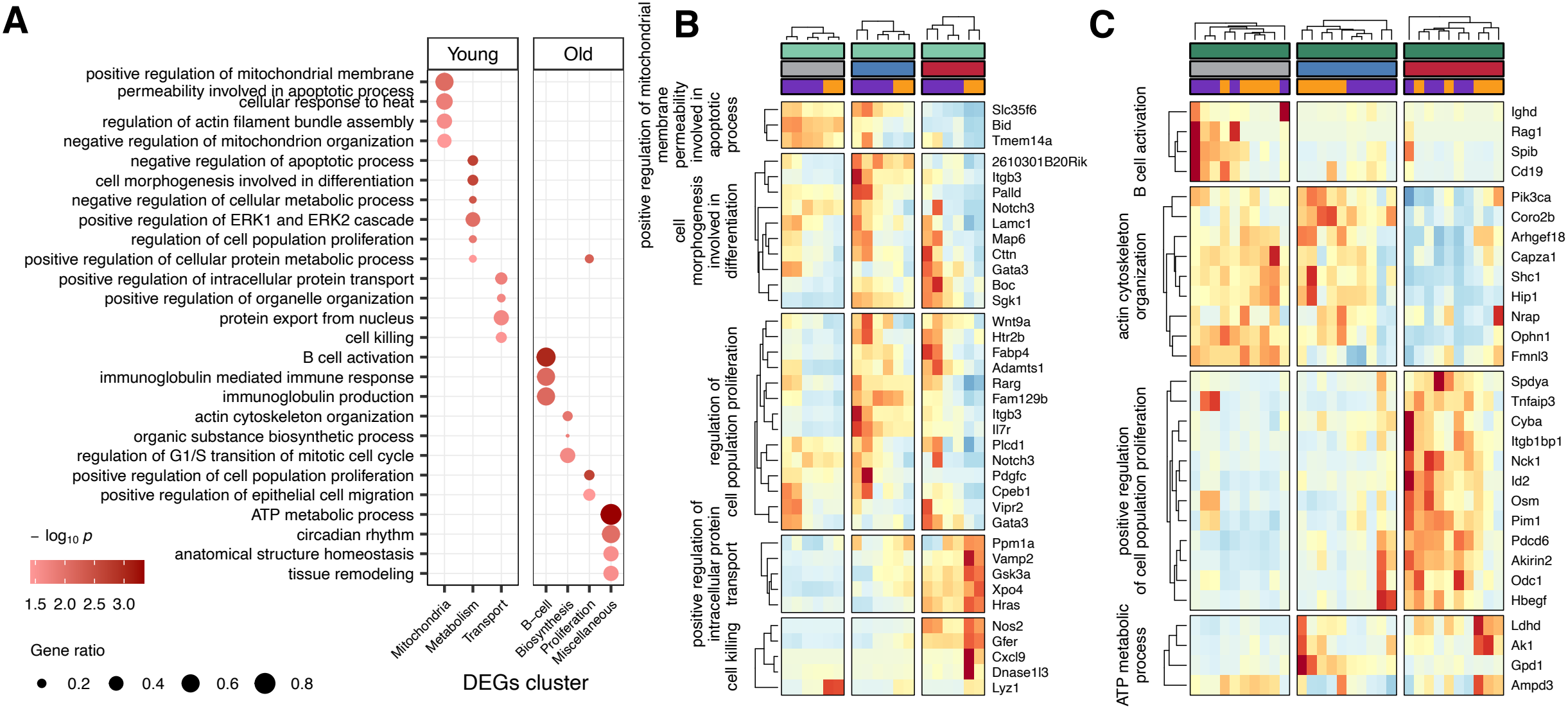
(A) Gene ontology biological process enriched in genes differentially expressed in polarized BMDMs from young and aged Dnmt3a^ΔLysM^ mice compared to Dnmt3a^fl/fl^ mice. Dot sizes indicate gene ratio as the radio of significant DEGs to annotated genes in each gene set term and colour corresponds to the p-value of enrichment. (B-C) Heatmaps showing scaled normalized expression of the leading DEGs between Dnmt3a^fl/fl^ and Dnmt3a^ΔLysM^ samples enriched in various biological processes. Analyses shown in A-C were conducted on n=4 young Dnmt3a^fl/fl^, n=2 young Dnmt3a^ΔLysM^, n=5 aged Dnmt3a^fl/fl^, and n=5 aged Dnmt3a^ΔLysM^ mice, with one BMDM sample per polarization per mouse; total n=48. Samples from young (8-10 weeks-old) or old (>34 weeks-old) mice are shown in light or dark green, respectively; while resting, M1 and M2 polarized BMDMs are shown in grey, red or blue, respectively; and wild-type (Dnmt3a^fl/fl^) or knockout (Dnmt3a^ΔLysM^) are shown in purple or orange. Heatmap columns are split according to hierarchical clustering and named according to key terms.

**Supplementary figure 3.**
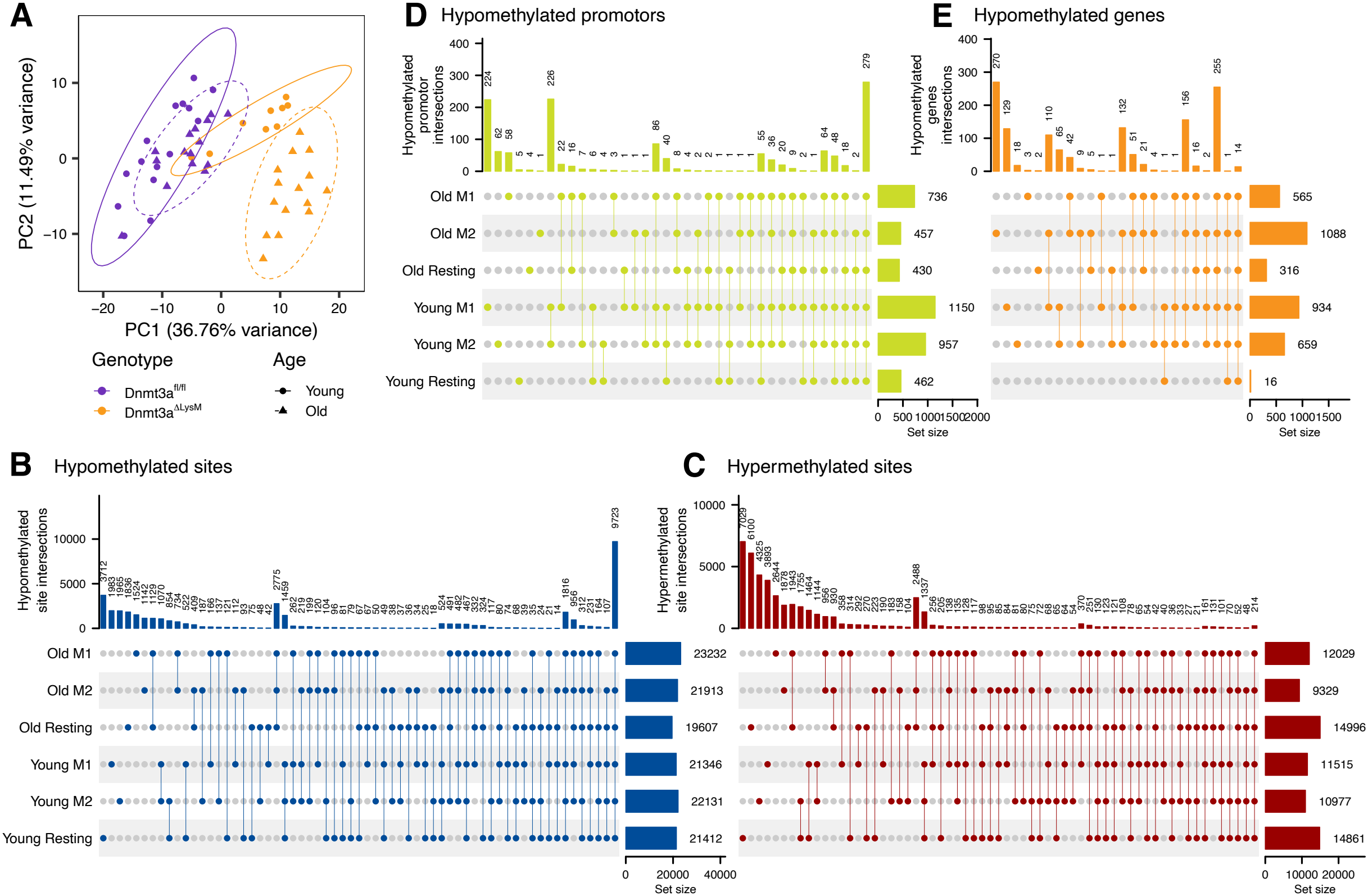
Dnmt3a loss induces hypomethylation in BMDMs. (A) Principal component analysis (PCA) of the DNA methylation array data for BMDMs from young (8-10 weeks old) and aged (>34 weeks-old) Dnmt3a^fl/fl^ and Dnmt3a^ΔLysM^ mice according to genotype and age. (B-C) Intersection plots of differentially methylated positions (DMP) in BMDM samples from young and aged Dnmt3a^fl/fl^ and Dnmt3a^ΔLysM^ mice according to hypomethylation (B) or hypermethylation (C). (D-E) Intersection plots of hypomethylated promoters (D) or genes (E) from young and aged Dnmt3a^fl/fl^ and Dnmt3a^ΔLysM^ BMDM samples. Analyses shown in A-E were conducted on n=5 young Dnmt3a^fl/fl^, n=3 young Dnmt3a^DLysM^, n=5 aged Dnmt3a^fl/fl^, and n=5 aged Dnmt3a^DLysM^ mice, with one BMDM sample per polarization per mouse; total n=54.

**Supplementary figure 4:**
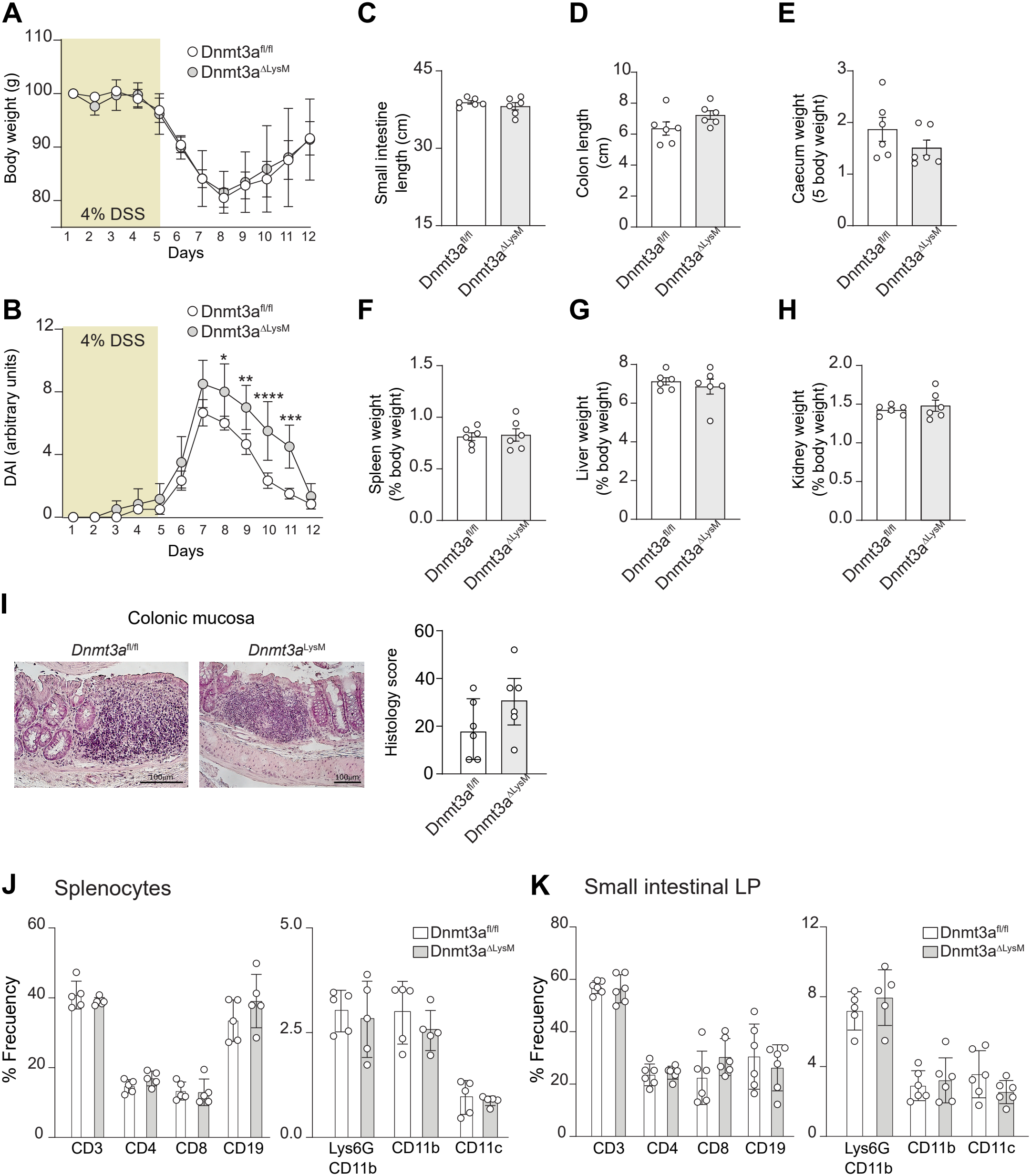
Dextran sodium sulfate (DSS)-induced colitis is stronger in mice lacking *Dnmt3a* expression in myeloid cells with lack of immune cell imbalance or organ macroscopic changes. (A-B) Percentage of body weight (A) and disease activity index (DAI, B) evolution during dextran sodium sulfate (DSS)-induced experimental colitis (4% applied on drinking water for 5 days as it is shown in brown) for Dnmt3a^fl/fl^ and Dnmt3a^DLysM^ mice. (C-H) Length of small intestine (C) and colon (D), and relative weights of caecum (E), spleen (F), liver (G) and kidney (H) at the completion of the experiment on day 12. (I) Representative images of colonic mucosa from Dnmt3a^fl/fl^ and Dnmt3a^DLysM^ mice after experimental colitis and histological scoring according to polymorphonuclear and mononuclear infiltration, crypt hyperplasia and epithelial erosion, and transmural inflammation. (J-K) Percentage of T-cells (CD3^+^), T helper cells (CD4^+^), cytotoxic T-cells (CD8^+^), B-cells (CD19^+^), neutrophils (Lys6G-CD11b^+^), macrophages (CD11b^+^) and dendritic cells (CD11c^+^) from spleen (J) or small intestinal lamina propria (K) from Dnmt3a^fl/fl^ and Dnmt3a^DLysM^ mice at the end of the experimental colitis experiment as assessed by spectral cytometry. All results are based on n = 5-6 mice per group. Data are represented as mean ± SEM.

**Supplementary figure 5:**
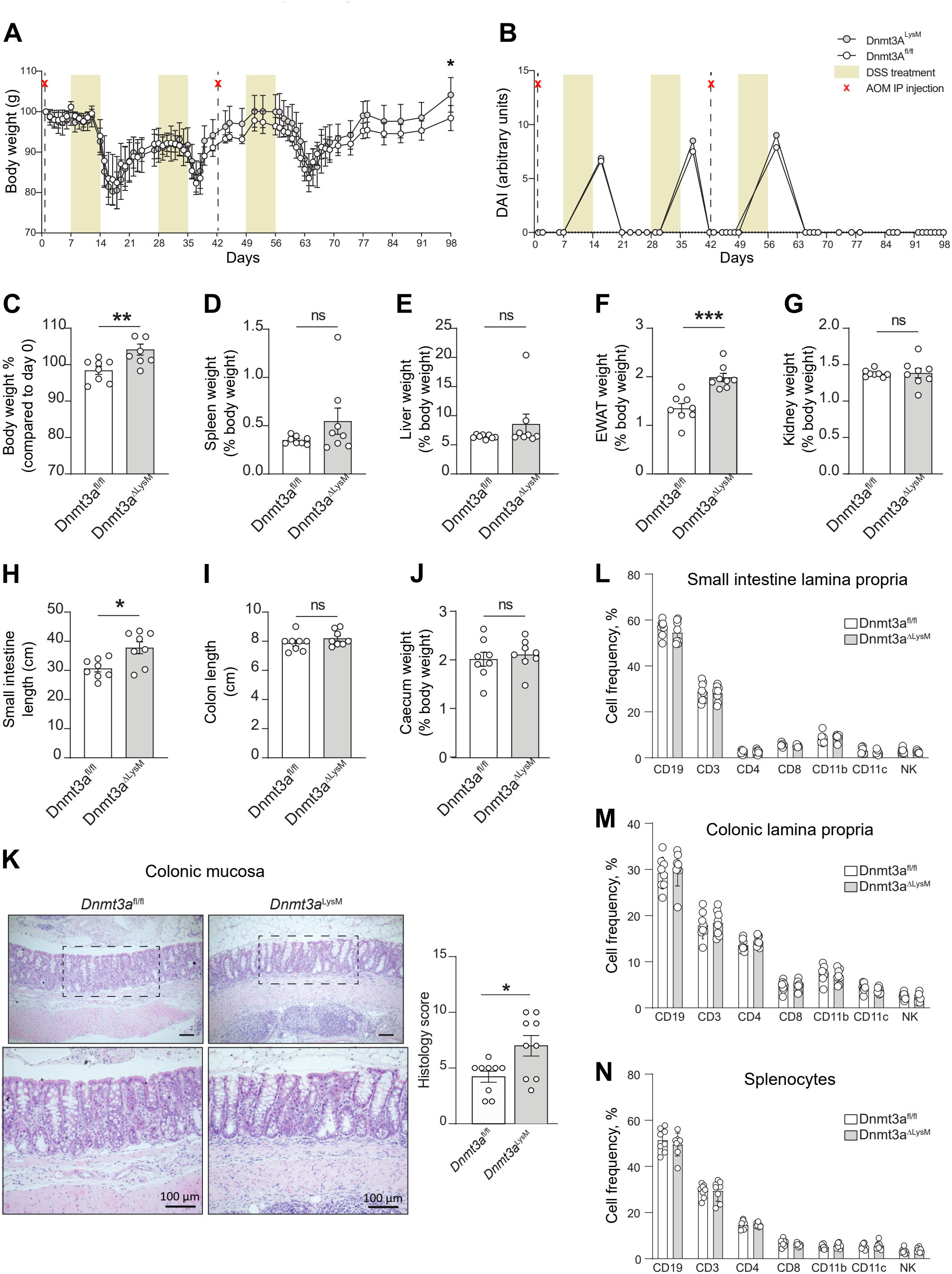
Azoxymethane (AOM)-dextran sodium sulfate (DSS)-induced chronic colitis induce tissue susceptibility to chronic inflammation without altering overall immune cell proportions in mice lacking *Dnmt3a*. (A-B) Body weight (A) and disease activity index (DAI, B) evolution during Azoxymethane (AOM)-dextran sodium sulfate (DSS)-induced experimental chronic colitis (1% DSS for cycle 1, and 1.5% DSS for cycles 2 and 3 on drinking water for 7 days as it is shown in brown with 10mg/kg AOM i.p. injections as it is shown by a red cross) for Dnmt3a^fl/fl^ and Dnmt3a^ΔLysM^ mice. (C-J) Percentage of body weight (C) and spleen (D), liver (E), epididymal white adipose tissue (EWAT, F) and kidney (G) according to body weight; length of small intestine (H) and colon (I), and relative weights of caecum (J) at the completion of the experiment on day 98. (K) Representative images of colonic mucosa from Dnmt3a^fl/fl^ and Dnmt3a^ΔLysM^ mice after experimental colitis and histological scoring according to polymorphonuclear and mononuclear infiltration, crypt hyperplasia and epithelial erosion, and transmural inflammation. (L-N) Percentage of different immune cells in the lamina propria of the small intestine (L) and colon (M), and cell frequencies among splenocytes (N) from Dnmt3a^fl/fl^ and Dnmt3a^ΔLysM^ mice at the end of the experimental colitis experiment as assessed by spectral cytometry. All results are based on n = 8 mice per group. Data are represented as mean ± SEM. Statistically significant differences are shown as *p<0,05, **p<0,001, ***p<0,0001.

